# Coordination of sequential RNase activities in an ancient molecular machine

**DOI:** 10.64898/2026.03.16.711897

**Authors:** Mathias Girbig, Francesca D. Naughton-Allen, Simone Prinz, Laura Andreas, Jan M. Schuller, Justin L. P. Benesch, Georg K. A. Hochberg

**Affiliations:** Evolutionary Biochemistry Group, Max Planck Institute for Terrestrial Microbiology, 35043 Marburg, Germany; Department of Biology, Philipps-University Marburg, 35043 Marburg, Germany; Department of Chemistry, University of Oxford, Oxford, UK; Kavli Institute for Nanoscience Discovery, Oxford, UK; Central Electron Microscopy Facility, Max Planck Institute of Biophysics, 60438 Frankfurt am Main, Germany; Center for Synthetic Microbiology (SYNMIKRO), Philipps-University Marburg, 35032 Marburg, Germany; Department of Chemistry, Philipps-University Marburg, 35032 Marburg, Germany

## Abstract

The evolutionary mechanisms by which molecular machines incorporate catalytic subunits and coordinate their functions are still poorly understood. Here, we study the molecular evolution of the RNA exosome, an essential RNA-processing machine built from a 9-subunit core (Exo9). Human and yeast Exo9 are catalytically inactive and serve as a recruiting hub for the peripheral RNase Rrp44. The core of modern exosomes descends from an active RNase in Archaea. Using ancestral sequence reconstruction, biochemical and structural characterization, we illuminate how Exo9 evolved from an enzyme to a regulatory hub. The ancestral Exo9 was an active, distributive RNase that already cooperated with Rrp44. Cryo-electron microscopy reveals how RNA-binding modulates the conformation of Exo9, thereby promoting allosteric recruitment of Rrp44. Exo9 begins substrate trimming before the RNA slips past its active site. The RNA can thereby be handed over to Rrp44 to be processed further, which rationalizes the coordination of consecutive RNase activities. We hypothesize that the same allosteric pathway still exists in the human exosome, implying that this mechanism persisted for over a billion years. Our work thus illuminates the evolutionary tinkering that produced a complex molecular machine and a central component of the eukaryotic cell.

## Main

Most molecular machines of eukaryotes descend from prokaryotic precursors that often have simpler topologies and fewer subunits. The mechanisms by which eukaryotic complexes evolved into their current forms are often poorly understood, because the histories of multi-subunit molecular machines and their activities are still hard to trace. One example of this is the eukaryotic RNA exosome. The RNA exosome degrades RNA molecules from 3’ to 5’ and carries out essential RNA processing and surveillance operations such as the trimming of ribosomal RNA^1^ and the decay of aberrant messenger RNAs^2^ (reviewed in Refs^3,4^). In humans and yeast, it consists of a catalytically inactive 9-subunit core (Exo9) forming a barrel through which RNA is threaded^5–7^. The barrel is bound by one or two peripheral RNases that contain the active sites of the complex: Rrp44 (also referred to as Dis3)^1,8^ and Rrp6^9^ (Fig. 1a). In yeast, they allosterically modulate each other^10^ binding Exo9 at opposing sites^11–13^. Both enzymes degrade RNA hydrolytically^8,14^ but Rrp6 is a distributive 3’-5’ exoribonuclease^5^, whereas Rrp44 is equipped with both endonucleolytic^15^ and processive exonucleolytic activities^8^.

**Fig. 1.**
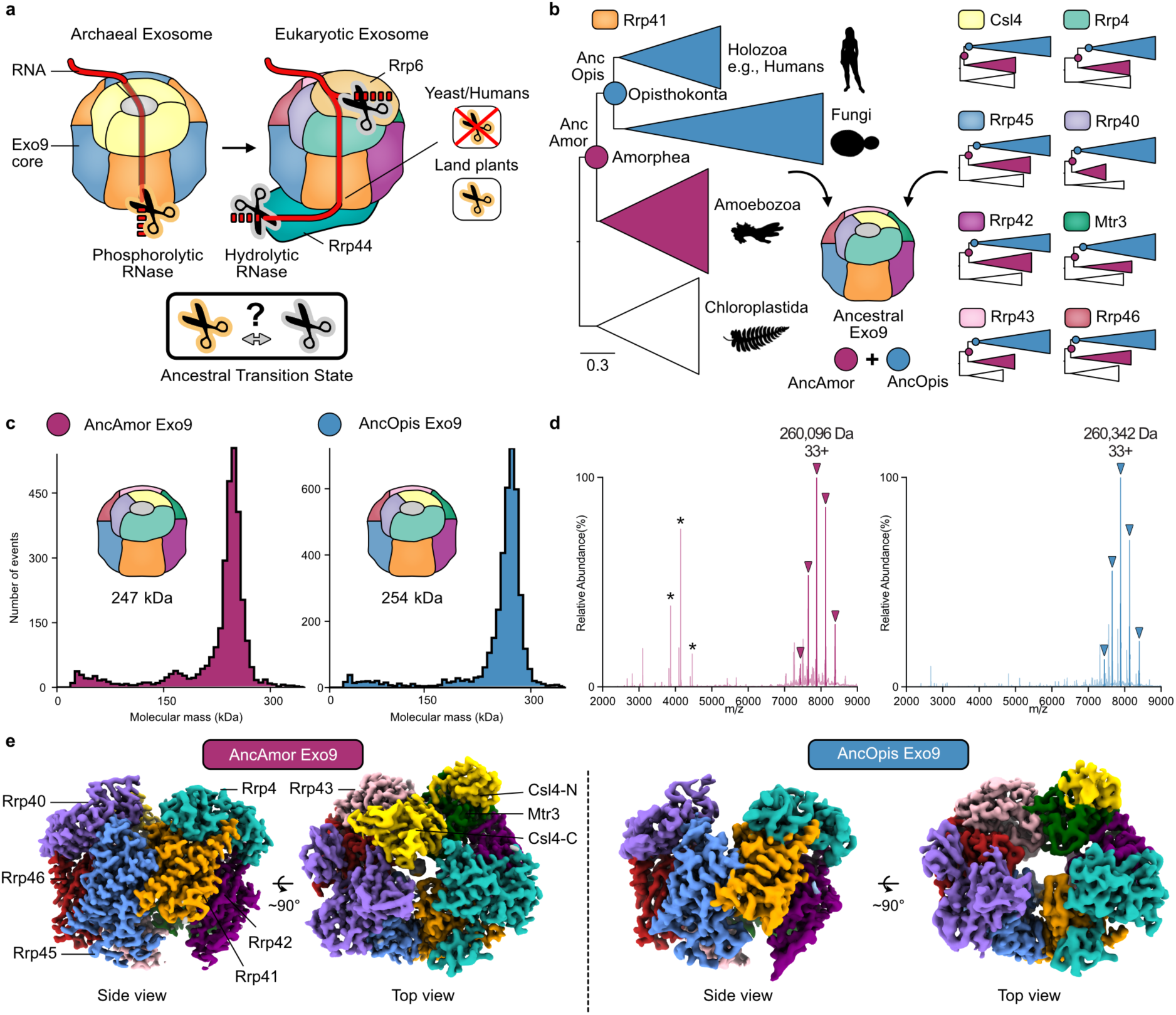
RNA exosome evolution and ancestral Exo9 resurrection. **a,** RNA exosome evolution and RNase activity in Archaea (left) and Eukaryotes (right). Evolution of eukaryotic RNase activity involved an ancestral transition state. **b,** Exo9 phylogenies and key nodes for ancestral Exo9 resurrection via ancestral sequence reconstruction (ASR). Left, reduced phylogeny for ASR of Exo9 subunit Rrp41 at indicated key nodes: Ancestral (Anc) Opisthokonta (Opis) and Amorphea (Amor). Silhouette images sourced from PhyloPic (http://phylopic.org). Scale bar: average substitutions per site. Right, reduced phylogenies of the other Exo9 subunits. **c,** Mass photometry (MP) of AncAmor (left) and AncOpis (right) Exo9 complexes. The measured molecular weights (MWs) of the main peaks are shown. The estimated MWs of successfully reconstituted Exo9s are 260.0 kDa (AncAmor) and 260.5 kDa (AncOpis). **d,** Native mass spectrometry of AncAmor (left) and AncOpis (right) Exo9 complexes. Colored arrows mark the protein species corresponding to correctly assembled Exo9 complexes. The measured MW of the most abundant complex in the specimens is shown. Asterisks mark a low abundance species: the Rrp41-45 heterodimer (Supplementary Table S2). **e,** Cryo electron microscopy (cryo-EM) structures of AncAmor (left) and AncOpis (right) Exo9 complexes. 3D reconstructions are shown from side and top views. Both complexes contain one copy per Exo9 subunit, as indicated with labels on AncAmor Exo9. The C-terminal domain of subunit Csl4 is not visible in AncOpis Exo9, indicating high mobility. Shown are cryo-EM maps, sharpened via EMready^43^.

Exo9 began its existence in Archaea as an active, processive RNase^16–21^ that worked independently of Rrp44 and Rrp6. The stoichiometry of the archaeal Exo9 is simpler, and it degrades RNA molecules using phosphate. How this processive RNase evolved to become a purely structural component and why it recruited peripheral RNases is not clear. The transition likely happened within Eukaryotes, because plants still retain an Exo9 core with phosphorolytic RNase activity^22^. Plant Exo9 also interacts with Rrp44, but it does so more transiently than its yeast counterpart^23–25^. This suggests that the evolution of the human and yeast exosome proceeded via an intermediate stage, in which an active Exo9 core was already coupled to a peripheral RNase. This would imply that the core activity was lost somewhere along the lineage to animals and fungi, transforming the exosome core from an RNase into a biochemical hub that organizes and recruits other RNases.

There are several conceptual difficulties with this trajectory. First, the plant Exo9 core is a distributive RNase^22^, unlike its processive archaeal ancestor. It is unclear if this activity reflects an ancestral state inherited by early eukaryotic exosomes from their archaeal ancestor, or whether it is a derived feature that evolved later, possibly along the lineage to plants. Further, the sequential arrangement of two active RNases is conceptually difficult to understand: Rrp44 binds the bottom of Exo9, and the RNA is first channeled through Exo9 before it reaches the exoribonucleolytic center of Rrp44. An active Exo9 core should therefore already degrade the RNA before it even reaches Rrp44’s active sites. If no RNA reaches Rrp44 via Exo9, what would be the purpose of their interaction? One possibility is that Rrp44 functions as a failsafe that rescues stalled RNA exosomes. Stalling could, for example, occur when the Exo9 core encounters structured RNAs, which it probably would not be able to degrade. But the active site of Exo9 is too deep within the barrel for Rrp44 to reach RNA that is stalled there. How Exo9 and Rrp44 functioned together in this ancient state, if indeed they did, and how their RNase activities were coordinated is therefore an unsolved problem.

These questions remain unanswered because the functional properties of ancestral molecular machines that existed deep in the history of eukaryotes are very difficult to infer from known properties of their extant descendants. This is because multi-subunit molecular machines are hard to purify and characterize. Experimental data are usually confined to a narrow set of genetically tractable model organisms, whose molecular machines may have undergone extensive lineage-specific evolution. Here, we overcome this challenge by combining ancestral sequence reconstruction (ASR)^26^ with biochemical, biophysical, and structural analysis of ancestral RNA exosome complexes that existed just before and just after the Exo9 core fully delegated its activity to Rrp44. Our results imply an ancient division of labor in the eukaryotic exosome, which had an active catalytic core that allosterically recruited Rrp44 upon RNA binding. This exosome takes off a few nucleotides before the RNA slips past the active site of the core and is further processed by Rrp44. This RNA-induced allosteric recruitment of Rrp44 is then inherited by RNA exosomes in which the core activity is fully lost. Our results rationalize the biochemistry of the human exosome as a consequence of its ancient evolutionary history and illuminate how the exosome core evolved from an active RNase to an inactive allosteric hub for other RNases.

### Reconstitution of resurrected ancestral Exo9 cores

To understand how the Exo9 core transitioned from an active RNase to an inactive hub for other RNases, we aimed to resurrect ancestral RNA exosome complexes from two major eukaryotic groups: Opisthokonta and Amorphea (Fig. 1b). Opisthokonta comprises fungi and animals, the only two groups that were experimentally proven to contain inactive exosome rings^5,6^. Amorphea represent a major eukaryotic supergroup and unite Opisthokonta with Amoebozoa. This clade was proposed to encode active Exo9 cores based on the conservation of residues necessary for RNase activity in the plant exosome^22^. We therefore hypothesized that the transition from an active to an inactive Exo9 core occurred on the branch leading from the last common ancestor of Amorphea (AncAmor) to Opisthokonta (AncOpis). To test this, we inferred phylogenies of the six RNA exosome ring subunits (Rrp41, Rrp45, Rrp46, Rrp43, Mtr3, Rrp42) and the three cap proteins (Csl4, Rrp4, Rrp40) using sequences from green algae and land plants as outgroups (Fig. 1b; Extended Data Figure 1). Because the inferred maximum likelihood (ML) phylogenies did not fully recapitulate the expected species topology within Amorphea (Extended Data Figure 1), we performed constrained tree searches, for which a set of clades (e.g., Holozoa, Fungi, Amoebozoa) were united whenever necessary. Approximately unbiased (AU) tests^27^ showed that none of the constrained trees was rejected by the data (Extended Data Figure 1), and we therefore proceeded with ASR using these trees.

For each Exo9 gene, we inferred the maximum a posteriori (MAP) ancestral protein sequences of the younger AncOpis, and the older AncAmor nodes, and recombinantly produced Exo9 proteins as described^28^ (Fig. 1b; Extended Data Figure 2 and Extended Data Figure 3a, b). We reconstituted the ancestral Exo9 complexes from AncOpis and AncAmor via size-exclusion chromatography (SEC) (Extended Data Figure 3c, d). Mass photometry (MP) analysis revealed that the ancestral Exo9 proteins assemble into homogenous complexes that are stable at nanomolar concentrations (Fig. 1c). Their measured molecular weights (MWs) closely matched the expected MWs (ca. 260 kDa) of Exo9 complexes containing a full complement of nine subunits. To examine the exact stoichiometry of these nonamers, we used native mass spectrometry (MS) and tandem MS/MS analysis (Fig. 1d; Extended Data Figure 4; Supplementary Table S1). For both AncOpis and AncAmor Exo9, native MS revealed that the most abundant complex for each was the 1:1:1:1:1:1:1:1:1 stoichiometry. We also measured a small fraction of sub-oligomeric species, but their abundance was much lower (Extended Data Figure 4; Supplementary Table S2). Hence, the vast majority of Exo9 proteins assembled correctly in ancestral Exo9 complexes.

To confirm that AncAmor and AncOpis Exo9 form functional RNA exosome complexes, we determined their cryo-electron microscopy (EM) structures at 2.7 and 3.6 Å resolution, respectively (Fig. 1e; Extended Data Figure 5 and Extended Data Figure 6; Supplementary Table S3). Both structures show the correct integration of all nine Exo9 subunits, and their architectures resemble those of fungal^29^ and human^5,30^ Exo9 complexes (Extended Data Figure 7). AncOpis and AncAmor Exo adopt similar structures overall, but we noticed that the C-terminus of Csl4 (Csl4-C) is not resolved in AncOpis, showing that it is more flexible than in AncAmor. A similar observation was made for the human Exo9 cryo-EM reconstruction^30^, indicating that flexible behavior of Csl4 is also present in modern RNA exosomes. Combined, our biophysical and structural analyses confirm the resurrection and reconstitution of ancestral RNA exosomes.

### Ancient exosomes had active and distributive Exo9 cores

Next, we tested the catalytic properties of the younger AncOpis Exo9 and the older AncAmor Exo9. To probe RNase activities, we added ancestral RNA exosomes to an AU-rich RNA oligonucleotide carrying a fluorescent tag at its 5’-end and monitored its length over time. AncAmor Exo9 robustly digested the RNA substrate in the presence of phosphate (Fig. 2a, lanes 2–5, Fig. 2b), whereas no RNase activity was observed for AncOpis Exo9 (lanes 7–10). When we omitted phosphate in the reaction buffer, AncAmor Exo9 did not show any signs of RNase activity (lanes 12–15). None of the AncAmor or AncOpis proteins could digest RNA by themselves (Fig. 2c, d, lanes 2–7). Only when the AncAmor proteins were mixed was the RNA substrate degraded (lane 8), confirming that AncAmor had an active Exo9 ring.

**Fig. 2.**
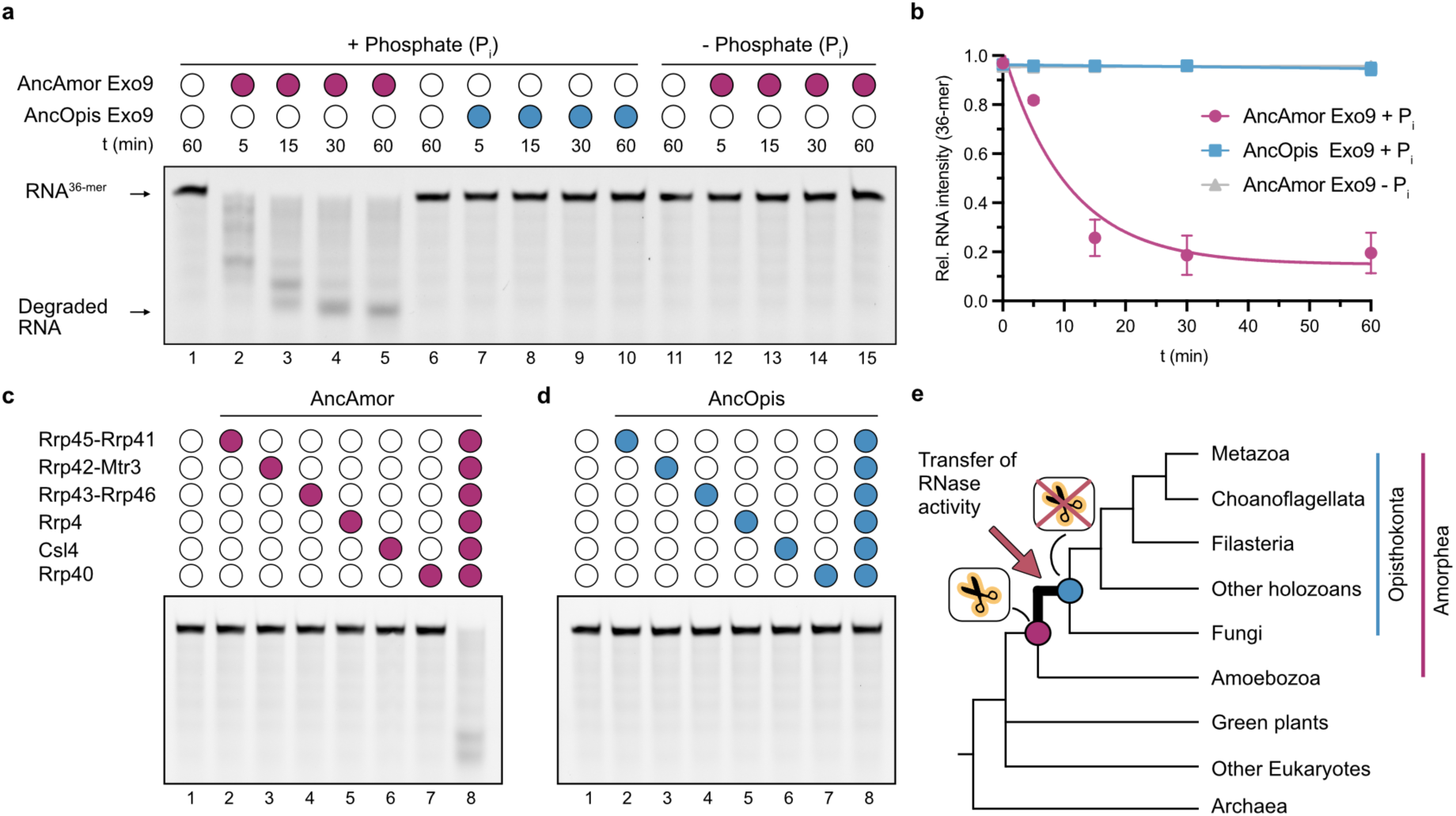
RNase activities of Ancestral Exo9. **a,** RNase assay of ancestral RNA exosomes in the presence (left) or absence (right) of inorganic phosphate. The RNA substrate carries a fluorescent carboxyfluorescein (FAM) label on the 5’-end. Shown is a representative denaturing acrylamide gel (15% urea, 1xTBE) of the three technical replicates. **b,** Quantification of RNase assays shown in panel (**a)**. Plotted is the intensity of full-length RNA (36-mer) compared to the total intensity of full-length and degraded RNA bands (mean values of three technical replicates (n = 3), error bars indicate standard deviation). Data points are fitted via a non-linear exponential decay model. **c–d** RNase assays of RNA exosome building blocks (monomeric cap and dimeric ring proteins) from AncAmor (**c**) and AncOpis (**d**). **e,** Simplified cladogram of selected eukaryotic groups, rooted with Archaea. Key nodes of interest are highlighted with colored spheres (purple: AncAmor, blue: AncOpis) and labeled with measured RNase activities of ancestral Exo9 cores. The red arrow marks the branch on which Exo9’s RNase activity got lost and transferred to the peripheral RNases during the history of Amorphea.

In both archaea and *A. thaliana* Exo9, subunit Rrp41 has been shown to harbor the catalytic residues to degrade the RNA phosphorolytically^17,22^. We therefore tested if the Rrp41−45 heterodimer is still sufficient for RNA degradation when implanted into an alternative exosome backbone. When we replaced AncOpis Rrp41−45 with its AncAmor counterpart, this hybrid Exo9 mix degraded RNA (Extended Data Figure 8a, lane 3). Vice versa, when we exchanged AncAmor Rrp41-45 with its AncOpis equivalent, RNase activity was lost (Extended Data Figure 8b, lane 2). Hence, the loss of RNase activity in AncOpis Exo9 is solely mediated through changes in Rrp41−45. We also tested if the inference on the ancestral Exo9 activities is robust to statistical uncertainty (Extended Data Figure 8c). We replaced Rrp41−45 with alternative, less likely (AltAll) reconstructions at both nodes while keeping the MAP ancestors of the other subunits. This confirmed that AncAmor Exo9 was catalytically active, whereas AncOpis Exo9 wasn’t (Extended Data Figure 8c). Thus, Exo9’s RNase activity was lost along the branch to Opisthokonta (Fig. 2e).

The RNase activities of both AncAmor and *A. thaliana* Exo9^22^ strongly suggest that the RNA exosome core of the last eukaryotic common ancestor (LECA) was still an active RNase, whose activity was only later lost. Just like the plant Exo9^22^, AncAmor Exo9 has distributive RNase activity, as indicated by the ladder-like RNA digestion pattern and late occurrence of the final RNA product band (Fig. 2a). This catalytic property differs from archaeal RNA exosomes, which are highly processive^18,19,21^. Thus, the transition from a processive to a distributive RNase likely occurred early during eukaryotic evolution, potentially even before LECA’s emergence.

### RNA-binding allosterically recruits Rrp44 to the active exosome core

We next asked if the ancestral Exo9 cores could already interact with the peripheral RNase Rrp44 from the AncOpis and AncAmor nodes (Fig. 3a) and resurrected ancestral AncAmor and AncOpis Rrp44 (Extended Data Figure 9a, b). Although the purified proteins showed minor proteolysis products (Extended Data Figure 9c), both enzymes efficiently degraded RNA, confirming the resurrection of functional RNases (Fig. 3b).

**Fig. 3.**
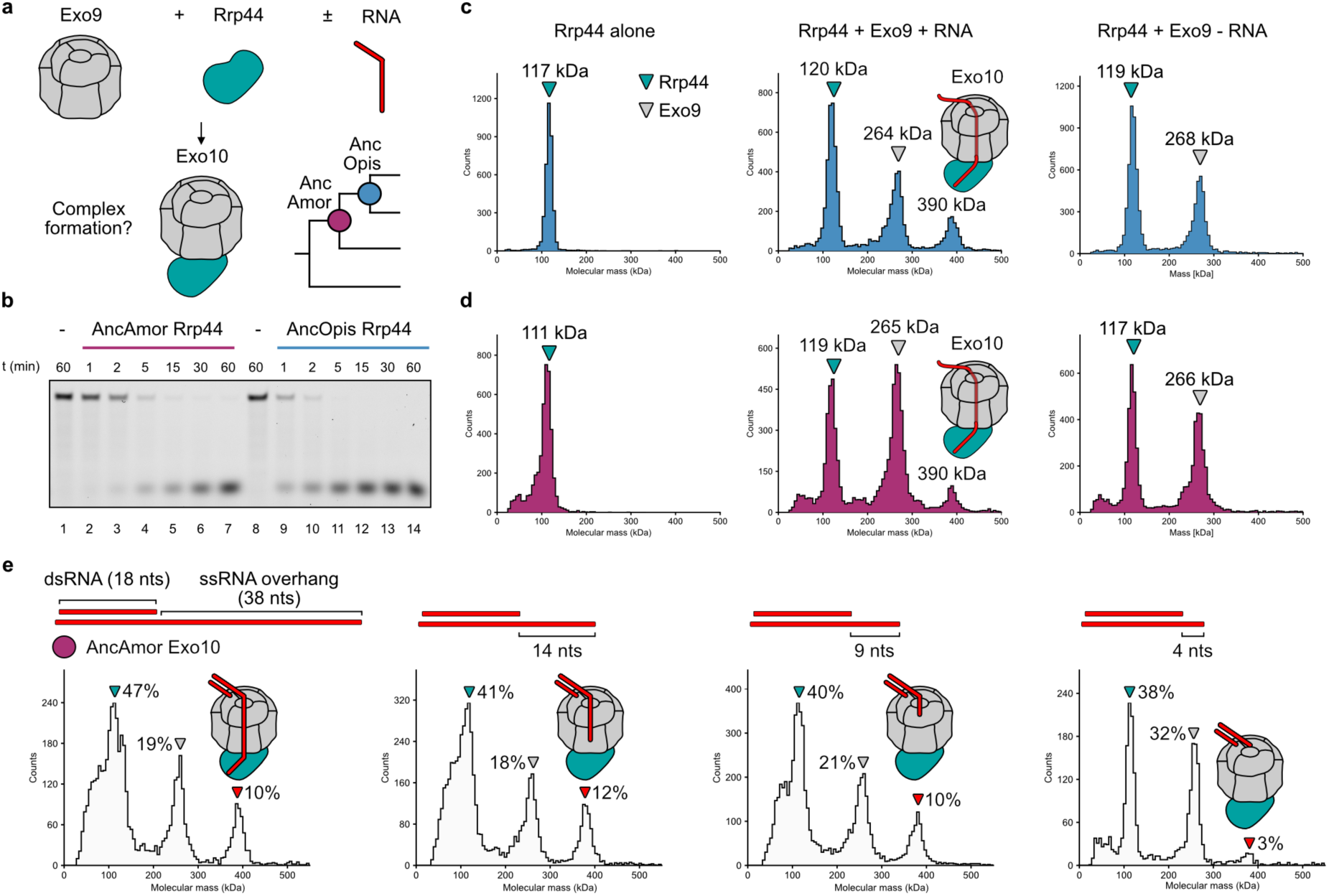
Allosteric recruitment of Rrp44 through RNA. **a,** Schematic of ancestral Exo10 complex formation out of Exo9 (grey) and Rrp44 (turquoise) tested in the absence or presence of RNA. Key nodes of interest are depicted in the cladogram on the bottom right. **b,** RNase assay of ancestral Rrp44 proteins, indicating that resurrected RNases are functional. **c–d,** Mass photometry (MP) analysis of AncOpis (**c**) and AncAmor (**d**) Exo10 components with measured molecular weights are given above the main peaks. Triangles mark Rrp44 (turquoise) or Exo9 (grey). Exo10 formation is only observed in the presence of RNA (middle panel). Exo10 peaks are highlighted with Exo10 schematics drawn above. **e,** MP analysis of AncAmor Exo10 complex formation in the presence of different RNA constructs reveals allosteric recruitment of Rrp44 to Exo9. All constructs contain the same 18 nucleotides (nts) long double-stranded (ds) RNA region and single-stranded (ss) RNA overhangs of different lengths (38–4 nts). Colored triangles (Exo9: grey, Rrp44: turquoise, Exo10: red) mark measured protein species. Percentage numbers indicate the relative abundance of molecular counts that were assigned to Rrp44, Exo9, or Exo10 species, with respect to the total counts in the mixture.

We used MP to test if the Exo9 cores and Rrp44 at the AncOpis and AncAmor nodes could form an Exo9+Rrp44 (Exo10) complex. AncOpis Rrp44 interacted with AncOpis Exo9 as shown by the appearance of a 390 kDa peak when they were mixed in the presence of RNA (Fig. 3c). This was expected as AncOpis Exo9 lost its activity and, therefore, fully relies on Rrp44 for processive substrate digestion. Interestingly, the older AncAmor Exo9 similarly interacted with AncAmor Rrp44 (Fig. 3d). Hence, AncAmor Exo9 already associated with Rrp44, even though AncAmor Exo9 was still an active RNase.

However, we did not observe a stable interaction between Exo9 and Rrp44 at both nodes when RNA was omitted (Fig. 3c, d). A potential explanation could be that the two molecules are linked through RNA, like beads on a string. Alternatively, the binding of RNA induces conformational changes within Exo9, thereby mediating allosteric recruitment of Rrp44. To test both hypotheses, we used a variety of RNA constructs harboring a double-stranded (ds) RNA region on the 5’-RNA end and a stretch of single-stranded (ss) RNA on the 3’-end. This construct design ensures that the RNA can only enter the Exo9 core from the top via its 3’-end (Fig. 3e).

We first analyzed a construct containing a long ssRNA overhang and observed robust Exo10 complex formation. We then gradually reduced the length of the ssRNA overhang to 14, 9, and 4 nts (Extended Data Figure 9d, e). Based on structures of RNA-bound exosome complexes^11–13,30^ and the experimentally determined footprint (ca. 20 nts) of the yeast Exo9 core^7^, these ssRNA stretches are too short to fully traverse the Exo9 core. Still, we observed equally efficient Exo10 formation with the 14 and 9 nt-long ssRNA overhangs (Fig. 3e). The association between Rrp44 and Exo9 must therefore be direct and induced by RNA-binding to Exo9. Even the shortest construct, containing only 4 nt-long ssRNA, shows signs of Exo10 complex formation, albeit at a lower efficiency. Using an alternative RNA construct that lacked a fluorescent FAM-tag and 3 nt-long ssRNA on the 5’-end while keeping the 14 nt-long ssRNA overhang on the 3’-end, we still observed Exo10 complex formation (Extended Data Figure 9f). Taken together, these results show that RNA binding in AncAmor recruits Rrp44 before the RNA emerges from the Exo9 core and probably even before it reaches AncAmor’s active site. Thus, the ancestral RNA exosome recruits Rrp44 through an allosteric mechanism, mediated by RNA binding.

### RNA-binding induces conformational changes that recruit Rrp44

To uncover the molecular basis of the RNA-mediated allosteric recruitment of Rrp44, we determined the cryo-EM structures of RNA-bound AncAmor Exo9 (Fig. 4a and Extended Data Figure 5d). The structural model of RNA-bound Exo9 was derived from the same datasets as used for the consensus AncAmor Exo9 structure because all Exo9 cryo-EM samples contained RNA. The consensus map of AncAmor Exo9 showed fuzzy density in the Exo9 center, suggesting that RNA is bound but also that it is mobile. Iterative rounds of 3D classification yielded a reconstruction of Exo9 bound to RNA (Extended Data Figure 5d) and an Exo9 map without RNA (Fig. 4b; Extended Data Figure 5e). Although the resolution was not sufficient to build an atomic model of the RNA, these two densities still allowed us to compare the conformations between RNA-bound and unbound Exo9 complexes (Fig. 4c). Whereas Csl4-C is completely unresolved in the absence of RNA, this region is well resolved in the RNA-bound Exo9 structure and interacts with the RNA (Fig. 4a).

**Fig. 4.**
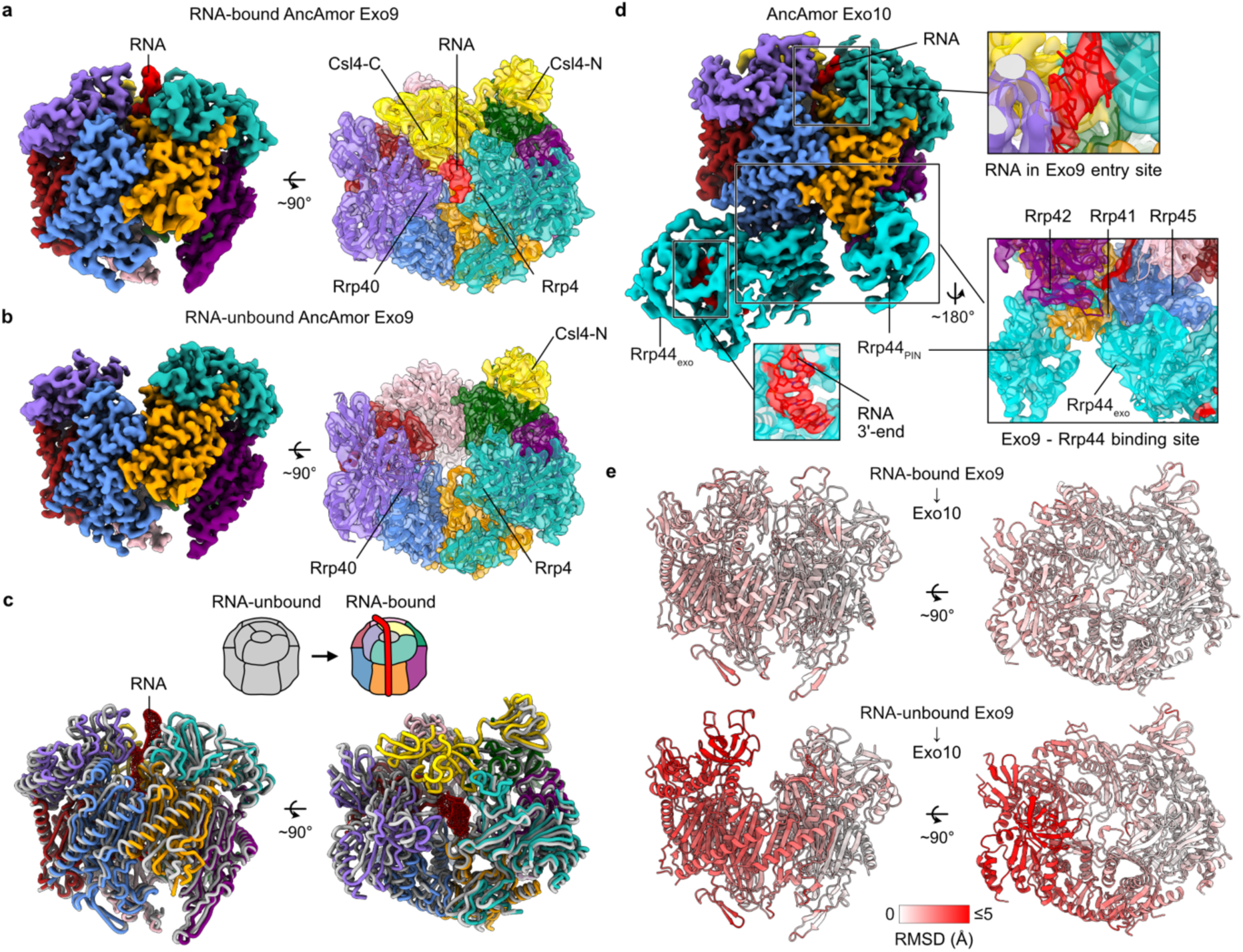
Structural basis of ancestral Exo10 complex formation. **a–b,** Cryo-EM maps of RNA-bound (**a**) and RNA-unbound (**b**) AncAmor Exo9 shown from the side (solid surface, left) and the top (transparent surface, right). The cryo-EM maps are colored according to the RNA exosome subunits taken from the structural models (see Fig. 1a for color code). The C-terminal region of subunit Csl4 (Csl4-C) is not resolved in the RNA-unbound cryo-EM map, indicating high mobility in the absence of RNA. **c,** Conformational changes upon RNA-binding. Structural models of RNA-unbound (grey) and RNA-bound (colored) are superimposed on subunit Rrp42 (purple). The RNA is depicted as a red mesh. **d,** Cryo-EM map of AncAmor Exo10 consisting of Exo9, RNA (red), and Rrp44 (turquoise) binding the bottom of Exo9. The N-terminal PIN and central exonuclease (Exo) domains are labeled. Close-up views indicate the presence of RNA at the top and center of Exo9 and in the active site of Rrp44 (bottom left). The resolution of the RNA-containing regions was not sufficient to build an atomic model of the RNA, but allows a putative fit of an RNA molecule in the Exo9 entry site (top close-up) and a phosphate backbone (bottom right close-up) in the center of Exo9. The binding site of Exo9 with Rrp44 is primarily formed by Rrp42, Rrp41, and Rrp45. **e,** The conformation of Exo10 resembles that of RNA-bound Exo9. Structures of RNA-bound Exo9 (top) and RNA-unbound Exo9 (bottom) are superimposed on AncAmor Exo10 (model not shown, superimposition on subunit Rrp42) and colored according to root mean square deviation (RMSD) between the superimposed models.

Superimposing the RNA-bound and unbound Exo9 structures reveals that RNA binding leads to contraction at the top of Exo9 and further rearrangements in the ring (Fig. 4c). We, therefore, asked if these structural rearrangements could mediate the recruitment of Rrp44 and determined the cryo-EM structure of AncAmor Exo10 at 3.3 – 3.6 Å resolution (Extended Data Figure 9g, h and Extended Data Figure 10). The 3D reconstruction resembled human RNA exosome structures^30,31^ and showed clear density for RNA and Rrp44 (Fig. 4d). To investigate how RNA binding permits Rrp44 binding, we superimposed the RNA-bound and unbound Exo9 structures onto AncAmor Exo10 and compared their root mean square deviation (RMSD) (Fig. 4e). The RMSD between AncAmor Exo10 and RNA-unbound Exo9 is much larger than it is to RNA-bound Exo9, suggesting that the interface of Exo9 in the absence of RNA is not compatible with the binding of Rrp44. Conversely, the structural rearrangement of Exo9 in the presence of RNA seems to favor the association with Rrp44. We therefore conclude that RNA-binding induces conformational changes in Exo9, which mediate the allosteric recruitment of Rrp44.

### Slippage facilitates RNA handover from Exo9 to Rrp44

Lastly, we addressed the questions of how the active Exo9 core functions together with Rrp44. We tested RNA degradation by AncAmor Exo9 and Exo10 on a 59 nt-long ssRNA with and without a dsRNA region at the 5’-end (Fig. 5a, b). Both complexes degraded the ssRNA substrate (Fig. 5a, b lanes 2-7), but we noticed a clear activity difference between Exo9 and Exo10. Exo9’s RNase activity is distributive, yields various intermediate bands, and only digests the substrate down to the final product length after ca. 120 min (Fig. 5a, lane 7). In contrast, Exo10 is processive and digests most of the RNA down to the final product already after 5 min without many intermediate products (Fig. 5b, lanes 2-7).

**Fig. 5.**
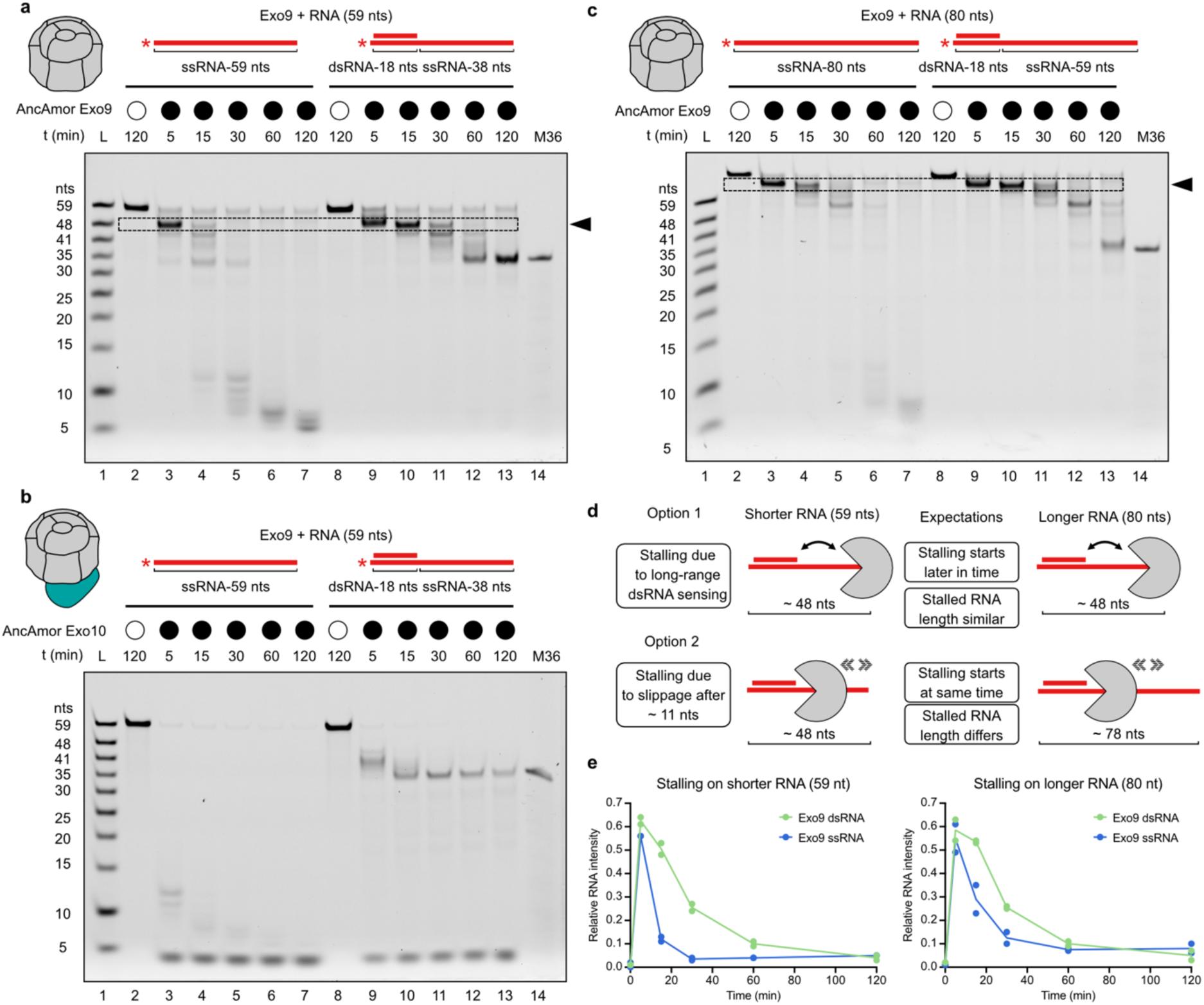
Rrp44 boosts RNA exosome activity and bypasses slippage-mediated Exo9 stalling. **a–c,** RNase assays of AncAmor exosomes with RNAs being fully single-stranded (ss) (lanes 2–7) or containing a double-stranded (ds) region (lanes 8-13) as indicated by the schematics shown above lanes. **a,** Exo9 on shorter RNA constructs with 59 nucleotides (nts). RNA degradation on the dsRNA construct is blocked at ca. 36 nts and stalled earlier after ca. 48 nts. **b,** Exo10 on shorter RNA constructs shows boosted RNase activity. Although dsRNA also hampers Exo10 activity, most of the substrate gets degraded over time. **c,** Exo9 on longer RNA constructs with 80 nts. Dashed boxes and black arrows indicate early stalling on shorter and longer dsRNA constructs. L: RNA ladder, M36: 36-nt long RNA marker. **d,** Schematics outlining potential explanations of Exo9 stalling on dsRNA constructs. Dashed arrows (option 1) indicate diffusive movement on RNA due to slippage, potentially resulting in early stalling. Two-headed arrows (option 2) depict potential long-range sensing of dsRNA, which could induce stalling. Experimental data favors option 1, considering the expectations for the different models. **e,** Quantification of relative intensity (selected band intensity divided by total lane intensity) of stalled RNA products (dashed boxes) observed for AncAmor Exo9 on shorter (**a**) and longer (**c**) RNA constructs. Green: Exo9 on dsRNA, blue: Exo on ssRNA. Plotted are data points from two technical replicates (n = 2). Mean values are connected by lines.

The effect on RNase activity is even more pronounced on the dsRNA substrate (Fig. 5a, b lanes 8-13). Exo9 trims the ssRNA overhang but completely fails to digest the dsRNA (Fig. 5a, lane 13). In contrast, Exo10 degrades the dsRNA substrate down to the final product length of ca. 4 nts (Fig. 5b, lanes 8-13). Although Exo10’s activity is also dampened by the dsRNA, the signal of the RNA hybrid still diminishes over time, whereas it remains stable in the presence of Exo9. Using an electrophoretic mobility shift assay (EMSA) and catalytically inactive Rrp44, we confirmed that the vast majority of Rrp44 associates with Exo9 under the experimental conditions (Extended Data Figure 9i). This indicates that the catalytic activity primarily stems from Rrp44, which is incorporated into Exo10 and not from free Rrp44 in the reaction mix. Thus, Rrp44 facilitates the degradation of RNAs with more complex secondary structures.

We also noticed a prominent RNA band produced by AncAmor Exo9 at earlier time points. This RNA species is more abundant on dsRNA when we tested the shorter construct with a 38 nt-long ssRNA overhang and a longer construct with a 59 nt-long overhang (Fig. 5a, c dashed frames). The signal persists for a longer time on dsRNA, indicating additional stalling on dsRNA substrates. The early stalling on a construct with dsRNA could have two explanations (Fig. 5d). The first option is that the dsRNA might be sensed by Exo9 through a long-range mechanism (e.g., due to the torque or stiffness of the dsRNA). According to this model, stalling should occur at a fixed distance from the hybrid on the 5’ end. It would begin later on the longer RNA because Exo9 takes more time to reach the critical stalling distance. The second option is that stalling occurs because the RNA slips past the active site after Exo9 digests a few nucleotides. Exo9 then might diffuse in either upstream or downstream direction and could only re-initiate RNA digestion once the 3’-end has slipped back into its active site or the RNA dissociates fully. According to this scenario, stalling on the longer RNA would be expected to occur at an RNA at the same distance from the 3’ end on the shorter and the longer RNA. To test both models, we compared Exo9’s RNase activity on the shorter and the longer dsRNA constructs and quantified the stalled RNA intensity over time (Fig. 5a, c, e).

We found that stalling already begins after 5 min at both constructs and persists in both for at least 30 min. Furthermore, the length of the stalled RNA substantially differed (shorter RNA: ca. 48 nts; longer RNA: ca. 60–70 nts). These results support the model that Exo9 stalls because of slippage and that the enzyme slips predictably after some 10 nucleotides. This explains how Rrp44 can engage with RNA that is first passed through the active Exo9 core: Rrp44 binds the RNA as soon as it slips past the active site and emerges from the pore of Exo9. Our data also provide a biochemical mechanism for why Exo9 is distributive: slippage naturally allows for RNAs of different lengths to be produced. Without a double-stranded hybrid on the 5’-end, RNA can probably slide through the exosome tunnel completely. This would explain why we see more stalling on substrates with a hybrid, where only one slide direction can re-engage the active site.

When acting together, Exo9 and Rrp44 do not stall (Fig. 5b) and quickly degrade the RNA without intermediates. This is also consistent with our MP data, which show that Rrp44 engages with the exosome even before it slips, as soon as RNA binds to Exo9. These results imply a programmed handover from AncAmor Exo9, which trims a few nucleotides phosphorolytically, then slips and passes the RNA on to an already primed Rrp44 at the exit from the Exo9 pore.

## Discussion

In this work, we illuminated a key step in the evolution of the RNA exosome. The ancestral state for this molecular machine in eukaryotes is to have an active, distributive Exo9 core, which allosterically recruits Rrp44 as soon as it binds RNA. The two RNases then function in sequence, which is only possible once Exo9 core slips after a small number of nucleotides. It may seem counterintuitive to incorporate slippage of an RNase on its substrate into the highly orchestrated mechanism of an essential molecular machine. But this is simply a dramatic example of tinkering – of using what works, regardless of its mechanistic elegance^32^.

Because Rrp44 is recruited immediately upon binding of RNA to the Exo9 core, it is very unlikely that Rrp44 is simply a failsafe mechanism for active Exo9 cores that stall on structured RNAs. Instead, the slippage-mediated handover mechanism we discovered here is likely used for precise trimming of certain RNAs. Based on data from the plant RNA exosome, the likely targets of this mechanism are RNA fragments that are produced during rRNA maturation^22^. We can only speculate about why the ancestral Amorphean exosome slips on RNA, whereas the processive Archaeal RNA exosome does not. One reason could be that the Archaeal exosome has three symmetrical active sites in its core^21^, whereas AncAmor has only one. This is due to the gene duplications of the exosome subunits that happened along the branch from Archaea to the last eukaryotic common ancestor. Accordingly, only the Exo9 subunit Rrp41 has retained RNase activity, and it is possible that this allows for slippage. Yet it is equally possible that the transition from a processive to a distributive RNase was driven solely by changes within the catalytic subunit Rrp41 itself, rather than by modifications in stoichiometry.

Yeast Rrp44 tightly binds Exo9 without RNA^5^, whereas human Rrp44 only transiently interacts with the exosome core^30^ – just like its ancestors did more than a billion years ago. The tighter binding to the exosome core by yeast Rrp44, on the other hand, represents a more derived, and potentially lineage-specific, arrangement. In agreement with this model, AncAmor Rrp44 adopts a conformation similar to that of human Rrp44^30,31^, resembling the ‘open conformation’ reported for some yeast RNA exosome complexes^12,13,33–35^. In contrast, the alternative, ‘closed conformation’ of yeast Rrp44^11,36,37^ is absent in AncAmor Exo10. The RNA also follows the same path as in human RNA exosome structures, which differs from that in yeast Exo9-Rrp44 complexes. Consequently, the structure of AncAmor Exo10 more closely resembles human Exo9-Rrp44 complexes than yeast ones. We therefore hypothesize that the allosteric recruitment of Rrp44, mediated through RNA-binding, persists in the human system.

The allosteric recruitment of Rrp44 has intriguing implications for how other components interact with the RNA exosome. Rrp6 is known to stimulate Rrp44’s activity when both are bound to opposite sides of Exo9^10^. Binding of Rrp6 also widens the PH ring at the top of the RNA exosome, similar to what we observe with RNA binding. We therefore suspect that this phenomenon uses the same allosteric mechanism as we discovered here. Future work will have to determine if Rrp6 already had this effect in exosomes that still have an active core. The RNA exosome also associates with a battery of cofactors, such as the RNA helicase MTR4^38^, and the TRAMP^39,40^, NEXT^41^, and PAXT^42^ complexes. The recruitment of Rrp44 through the RNA substrate itself suggests that Rrp44 integration is modular. This modularity may allow Rrp44 to hop from one exosome to another, potentially depending on which auxiliary factors these exosomes interact with. In this regard, it will be interesting to determine whether and how these auxiliary factors differentially modulate Rrp44 recruitment. This, in turn, may depend not only on the nature of the RNA substrate but also on the organism and lineage to which it belongs. Indeed, the stable integration of Rrp44 into the RNA exosome of baker’s yeast, in contrast to its transient association in both extant and ancestral exosomes, already shows that this is the case. We therefore anticipate that evolution has not only shaped the activity of the RNA exosome core but also its function as an allosteric hub and its interplay with cofactors.

Our study demonstrates that entire molecular machines can be resurrected and that the properties of these ancient machines have much to tell us about why their modern descendants look and function the way that they do.

## Methods

### Phylogenetic analysis and ancestral sequence reconstruction

Homologous protein sequences of exosome subunits from species covering the clades Opisthokonta, Amoebozoa, and Chloroplastida were retrieved via BLAST^46^ from the EukProt v3 database^47^. The database was downloaded, and the BLAST searches were carried out locally using Sequenceserver^48^ and the human exosome subunits as query sequences. Sequences were aligned using MAFFT^49^ or MAFFT-DASH^50^, and the multiple sequence alignments (MSAs) were inspected and trimmed manually using Aliview^51^. If necessary, profile alignments of orthologous sequences belonging to the same clade were first generated, processed as described above, and afterwards combined using the profile-align function in MUSCLE^52^.

Maximum-likelihood (ML) phylogenetic trees were computed using IQ-TREE^53^. Substitution models were selected using ModelFinder^54^, and statistical branch support values were obtained via the ultrafast bootstrap approach^55^ and the SH-like approximate likelihood ratio test (1000 replicates)^56^. ML tree searches were repeated at least three times, and the trees with the best likelihood were chosen for subsequent analysis. Inferred phylogenies of individual exosome components were inspected via FigTree^57^ and rooted using the orthologous proteins from chloroplastida as an outgroup. In the case of deviation between a gene tree and the amorphean species tree at the nodes of interest (e.g., AncOpisthokonta), constrained tree searches were performed. Multifurcating trees were generated via the Mesquite^58^ software to follow the amorphean species topology at deeper nodes and used as guide trees for constrained tree searches in IQ-TREE. The ML trees and computed constrained trees were subjected to approximately unbiased (AU) tests^27^ with 10,000 replicates per test, which did not reject any of the tested constrained trees.

For ancestral sequence reconstruction (ASR), the rooted constrained trees of the individual exosome core subunits and the ML tree of Rrp44 were used. Ancestral gaps were assigned via PastML^59^, and ancestral sequences were inferred with IQ-TREE by the empirical Bayesian method^60^. This estimates the posterior distribution of states per site given the provided MSA, the tree, and model parameters. The maximum a posteriori (MAP) sequences, which contained the states with the highest posterior probabilities (PP) at each site, were inferred. The alternative ancestral amino acid sequences (of Rrp45, Rrp41, Rrp44) were also inferred, which contained the states with the second-best PPs per site in case it was > 0.2.

### Molecular cloning

Protein-coding genes were codon-optimized for protein production in *Escherichia coli* and ordered as gene fragments from Twist Bioscience. Ancestral exosome proteins and those from extant organisms were, except for Csl4 proteins, designed to have an N-terminal His_6_-tag, followed by a tobacco etch virus protease cleavage site (TEV). Csl4-coding genes contained an N-terminal Twin-Strep tag, followed by an N-terminal His_6_-tag and a human rhinovirus (HRV) 3C protease site. The ancestral amorphean protein-coding genes were ordered together with upstream T7 promoter- and downstream T7 terminator sequences and flanking BsaI recognition sites. The fragments were cloned via BsaI (BsaI-HFv2, NEB) into a custom version of the pCDF vector, engineered to be applicable for Golden Gate cloning. For all other genes, which were cloned at later stages during this project, gene fragments were ordered as open reading frames, flanked by SapI recognition sites. The fragments were cloned into custom pCDF vectors, which were engineered to carry either no affinity tag (used for Rrp45, AncOpis Mtr3, Rrp43), an N-terminal His_6_-TEV-tag (used for Rrp41, AncAmor Mtr3, Rrp46, Rrp40, Rrp4, Rrp44), or an N-terminal Twin-Strep-His_6_-HRV3C (used for Csl4). The vectors were designed to allow scarless, in frame-cloning^61^ of the genes via SapI (NEB) using Golden Gate cloning. The exosome ring genes were further cloned for co-expression as pairs (Rrp45-Rrp41, Rrp42-Mtr3, Rrp43-Rrp46). The full gene expression cassettes (each gene with its own promoter and terminator) of the respective pairs were joined together and cloned via BsaI (NEB) into a custom pCDF vector (for the AncAmor ring proteins) or pRSF vector (for all other ring proteins). Cloning was performed using chemically competent *E. coli* DH5-alpha cells and standard molecular cloning practices. Site-directed mutagenesis was performed via the two Single-Primer Reactions IN Parallel (SPRINP) method^62^, for which Phusion™ High-Fidelity DNA Polymerase (NEB) was used to generate linear single-primer PCR products. Spectinomycin (100 µg/mL) was used for pCDF-derived constructs, and kanamycin (50 µg/mL) was used for pRSF-derived constructs. All cloned constructs were sequenced either via Sanger sequencing or full plasmid sequencing at Microsynth Seqlab GmbH.

### Protein production

Expression and purification procedures of ancestral exosome proteins (AncAmor and AncOpis) were designed based on published experimental strategies to produce yeast or human exosome proteins^28^. Exosome cap proteins (Rrp4, Rrp40, Csl4) and Rrp44 variants were produced separately using single-gene expression constructs. Exosome ring proteins were co-expressed as pairs (Rrp45-Rrp41, Rrp42-Mtr3, Rrp43-Rrp46). For Rrp45-Rrp41 and Rrp43-Rrp46 (both AncAmor and AncOpis), and Rrp42-Mtr3 (AncOpis), the expression constructs were designed so that only one (Rrp41, Mtr3, Rrp46) of the protein pairs carried an N-terminal His_6_-tag. For AncAmor Rrp42-Mtr3, two expression constructs were used (His_6_Rrp42-Mtr3 and Rrp42-His_6_Mtr3). Expression constructs were transformed into chemically competent *E. coli* BL21 (DE3) cells via heat-shock and plated onto Lysogeny Broth (LB) plates. LB plates and all liquid expression media contained the required antibiotics for the pCDF or pRSF constructs (see above). Overnight cultures were inoculated either from single colonies or glycerol stocks. The next day, the overnight cultures were used to inoculate expression cultures (typically 0.5 to 2 L), for which Super Broth (SB) media (3.5% tryptone, 2% yeast extract, 0.5% NaCl, pH 7.5) was used. Cells were grown at 37 °C and 240 rpm to an optical density of 1.0 to 1.5 and kept on ice for at least 10 minutes. Protein gene expression was induced by adding isopropyl β-D-1-thiogalactopyranoside (IPTG) to a final concentration of 0.5 mM, and the cultures were incubated at 18°C and 140 rpm overnight. The next day, cultures were harvested through centrifugation (5000 rpm, F9-6×1000 LEX rotor (ThermoScientific), 15 min at 4 °C), and the cell pellets were resuspended in 1 mL of resuspension buffer (50 mM Tris-HCl, pH 8.0, 20% sucrose) per 0.5 g of cell pellet and kept at −20 °C until usage. For AncAmor Rrp42-Mtr3 samples, cultures that separately expressed His_6_Rrp42-Mtr3 and Rrp42-His_6_Mtr3 were mixed and harvested together. The resuspension buffer and all other Tris-HCl-based buffers were pH-adjusted at room temperature.

For protein purification, cell suspensions were thawed in a room-temperature warm water bath and, once thawed, kept on ice for further usage. Cell suspensions were supplemented with an equal volume of lysis buffer (50 mM Tris-HCl, pH 8.0, 500 mM NaCl, 20 mM Imidazole, 20% sucrose, 0.1% IGEPAL, 1 mM β-mercaptoethanol (BME), DNase (10 µg/mL), 1 cOmplete Protease Inhibitor Cocktail (Roche) (1 tablet per 100 mL)). Cells were lysed on ice via sonication (70% amplitude, 30 s pulse, 90 s pause, 3.5–4.5 min total) using an SFX150 digital sonifier (Branson), equipped with a 3.2 mm microtip. Cell lysates were cleared via centrifugation (15.900 rpm, Sorvall SL-50T rotor (ThermoScientific), 30 min at 4 °C), the supernatants were filtered using 0.45 µm syringe filters (Sarstedt), and subjected to immobilized metal affinity chromatography (IMAC) using either 5 mL Nuvia IMAC Ni-charged (Bio-Rad) or 5 mL HisTrap HP (Cytiva) columns, equilibrated in IMAC wash buffer (10 mM Tris-HCl, pH 8.0, 350 mM NaCl, 20 mM Imidazole, 1 mM BME). The columns were loaded for 30–40 min using a peristaltic pump (LabN1-III (Shenchen) or Hei-FLOW Precision 06 (Heidolph)) and washed with at least 10 column volumes (CV) of IMAC wash buffer (WB). All subsequent chromatography steps were performed at 4 °C, using an NGC Chromatography System (Bio-Rad). Peak fractions were analysed via SDS–polyacrylamide gel electrophoresis (PAGE) and fractions containing the proteins of interest were pooled. For subsequent protein concentration steps, Ultra Centrifugal Filters (Amicon, Merck) were used, which were also used for buffer exchange. Protein concentrations were determined using UV absorption at 280 nm and construct-specific extinction coefficients. To elute nickel-bound proteins from the IMAC columns, a gradient from IMAC WB to IMAC elution buffer (EB) (10 mM Tris-HCl, pH 8.0, 350 mM NaCl, 500 mM Imidazole) was applied. Subsequent chromatography steps differ between proteins as described below.

Exosome cap proteins (Rrp4, Rrp40, Csl4) were concentrated and subjected to size-exclusion chromatography (SEC) using either an ENrich SEC 650 column (Bio-Rad) or a HiLoad 16/600 Superdex 200 pg column (Cytiva), equilibrated in SEC350 buffer (20 mM Tris-HCl, pH 8.0, 350 mM NaCl, 1 mM Dithiothreitol (DTT)). Peak fractions were concentrated to typically 6– 13 mg/mL (Rrp4), 3–6 mg/mL (Csl4), and 30 mg/mL (Rrp40), snap-frozen in liquid nitrogen and stored at −70 °C until usage.

Following IMAC, His_6_-tagged exosome ring protein pairs (Rrp45-His_6_Rrp41, His_6_Mtr3-His_6_Rrp42, and Rrp43-His_6_Rrp46) were subjected to overnight TEV cleavage at 4 °C, for which the NaCl concentration was adjusted to 150 mM using a mix of typically 20 mL ion exchange (IEX) buffer A (20 mM Tris-HCl, pH 8.0, 50 mM NaCl, 1 mM DTT) and 10 mL SEC150 buffer (20 mM Tris-HCl, pH 8.0, 150 mM NaCl, 1 mM DTT). The next day, the protein samples were concentrated, diluted with IEX buffer A to adjust the NaCl concentration to less than 100 mM, filtered with 0.22 µm syringe filters (Sarstedt), and subjected to a UNO Q1 column (Bio-Rad), equilibrated in IEX buffer A. The column was washed with IEX buffer A, and bound proteins were eluted using a linear gradient from 0–100% IEX buffer B (20 mM Tris-HCl, pH 8.0, 1 M NaCl, 1 mM DTT) over 35 mL. In the case of AncAmor and AncOpis Rrp45-Rrp41, AncOpis Rrp42-Mtr3, and AncAmor and AncOpis Rrp43-Rrp46, the eluted proteins of interest were buffer-exchanged into SEC350 buffer, concentrated, snap-frozen in liquid nitrogen, and stored at −70°C until usage.

For the purification of AncAmor Rrp42-Mtr3, the TEV-cleaved protein sample was subjected to a 5 mL HiTrap™ Heparin HP (Cytiva) column, using the same IEX buffers as for the UnoQ column. Eluted peak fractions were analyzed via a ribonuclease assay (see below), and those fractions with minimal RNase activity were pooled, treated as above, and then applied to the UnoQ1 column. Eluted peak fractions were also subjected to a ribonuclease assay, and those fractions with minimal RNase activity were pooled, concentrated, and applied to SEC in SEC350 buffer. SEC elution fractions were analysed once more with a ribonuclease assay, and only those fractions with no detectable RNase activity were pooled, concentrated, snap frozen, and stored at −70 °C until usage.

For Rrp44 proteins, IMAC-eluted samples were directly diluted using IEX A and applied to the Uno Q1 column. Bound proteins were eluted using a narrow linear gradient from 0–50% IEX buffer B over 50 mL. All eluted protein fractions of interest were analysed by SDS-PAGE, and those fractions that showed the least amount of degradation products (corresponding to the early eluting fractions) were pooled, concentrated, and subjected to SEC using SEC150 buffer. SEC peak fractions were pooled, concentrated, snap frozen, and stored at −70°C until usage.

### RNA exosome reconstitution

Ancestral Exo9 complexes were reconstituted by mixing equimolar ratios (50 µM each, input concentration) of the AncAmor or AncOpis ring (Rrp45-Rrp41, Rrp42-Mtr3, Rrp43-Rrp46) and cap proteins (Csl4, Rrp4, Rrp40). The mixture was filled up to a volume of 500 µL with SEC350 buffer to yield a concentration of 8 µM of the complex and dialyzed against 1L of SEC100 buffer (20 mM Tris-HCl, pH 8.0, 100 mM NaCl, 1 mM DTT) overnight at 4 °C using Mini Dialysis Kits (1 kDa molecular weight cutoff, Cytiva). The next day, the dialysis buffer was replaced with 1 L of fresh SEC100 buffer and dialysed for another 2 h. The samples were transferred to a fresh tube, centrifuged for 6 min at 14,000 g, and the supernatant was applied to an ENrich SEC 650 10 x 300 column (Bio-Rad), equilibrated in SEC100 buffer. Peak fractions were analyzed via SDS-PAGE and mass photometry (MP), and fractions containing fully assembled Exo9 complexes were pooled, concentrated to 10–40 µM using 0.5 mL ultracentrifugation filters (Amicon, 100 kDa molecular weight cutoff, Merck), snap frozen, and stored at −70°C until usage.

Ancestral Exo10 complexes were reconstituted by mixing equimolar amounts (8 µM each, final concentration in the mix) of catalytically inactive ancestral Rrp44, Exo9, and FAM-labelled ssRNA59 in SEC100 buffer supplemented with 5 mM MgCl_2_ in a total volume of 40 µL. The samples were incubated for at least 15 min on ice and applied to a Superose 6 Increase 3.2/300 column (Cytiva), equilibrated in SEC100 buffer. Peak fractions were analyzed via SDS-PAGE, native PAGE (see below), and MP. Fractions containing fully assembled Exo10 complexes were pooled and concentrated using 0.5 mL ultracentrifugation filters (Amicon, 100 kDa molecular weight cutoff, Merck) to a concentration of approximately 1.9 µM. The concentration was estimated using UV absorption at 280 nm and the expected extinction coefficient of the Exo10 proteins without considering the absorption of the RNA oligonucleotide. The complexes were snap frozen and stored at −70 °C until usage.

### Mass photometry

Mass photometry (MP) data of AncAmor and AncOpis Exo9 was acquired on a TwoMP mass photometer (Refeyn Ltd, Oxford) using AcquireMP software (V2.5.0, Refeyn Ltd.). Coverslips (Marienfeld, 22 mm x 50 mm #1.5) were cleaned three times with isopropanol and Milli-Q water and dried via a compressed air stream. Measurements were performed in silicon gaskets (3 mm x 1 mm, GBL103250, Grace Bio-Labs) attached to clean coverslips. Before the addition of the sample, SEC100 buffer was added to the gasket, and the focus position was adjusted to optimize the contrast at the glass-solution interface. For each measurement, sample was added to the gasket containing SEC100 buffer to a final concentration of typically 25 nM before a movie was recorded for 60 s using a frame rate of 48 frames per second (fps). Movie processing and particle detection were performed using DiscoverMP software (V2024 R1, Refeyn Ltd.). A custom molecular weight standard (84–336 kDa) was used for contrast-to-mass calibration.

For experiments probing the interactions between AncAmor or AncOpis Exo9 with Rrp44 (Fig. 3b,c), active versions of Rrp44 were used. For experiments using different lengths of ssRNA overhangs (Fig. 3e), inactive versions of Rrp44, for which the residues for endo- and exonucleolytic activity were mutated according to Ref^15^, were used. Inactive Rrp44 was used to allow longer incubation with the RNA and to ensure that recorded differences only stem from differences in RNA length and not from degradation activities by Rrp44.

For the Exo9 and Rrp44 interaction analysis, Exo9 and Rrp44 were pre-mixed in SEC100 at a 1:2 ratio and kept at room temperature. For each measurement, 3 µL were taken from the mix and freshly supplemented with either 1 µL SEC100 buffer (without RNA samples) or 1 µL of RNA (at 4 µM, prepared as described below). The mix was incubated at 37°C for 5 minutes, 0.5 µL was applied to 19.5 µL of SEC100 buffer, pre-added to the gasket, and the recording was started. The final concentrations in the drop were: Exo9 (12.5 nM), Rrp44 (25 nM), RNA (25 nM). For MP measurements of reconstituted Exo10 complexes, 0.5–1 µL of sample was taken directly from the collected SEC fractions, and measurements were acquired at 50 fps.

### Native and tandem mass spectrometry

Samples were buffer exchanged into 500 mM ammonium acetate (pH 6.9) using ultracentrifugation filters (Amicon, 10 kDa molecular weight cutoff, Merck) and diluted immediately before measurement using LC-MS grade water to a final concentration of 1.2 µM (AncAmor Exo9) or 2.0 µM (AncOpis Exo9) in 250 mM ammonium acetate (pH 6.9).

Nanospray capillaries were pulled in-house using a P97 Micropipette Puller (Sutter Instrument Corporation) and gold-coated using an Agar Auto Sputter Coater^63^.

Samples were delivered using a nano-electrospray ionization source in positive mode into a Q Exactive UHMR Orbitrap mass spectrometer (Thermo Fisher Scientific). Instrument parameters were tuned using Xcalibur software (Thermo Fisher Scientific) to maximize ion desolvation without fragmentation. The instrument parameters were maintained as follows: desolvation potential −140 V (AncAmor Exo9) or −130 V (AncOpis Exo9), trapping gas pressure 6.0, capillary temperature 150 °C, capillary potential 1.3 kV (AncAmor Exo9) or 1.1 kV (AncOpis Exo9), collisional activation in the HCD cell 0 V, and orbitrap resolution 12,500.

To acquire tandem MS spectra, an ensemble of ions within a given m/z window was selected using the quadrupole and subjected to collisional activation in the HCD cell of 120 V (AncAmor Exo9) or 115 V (AncOpis Exo9). Mass deconvolution was performed using UniDec version 6.0.3^64^ and MassLynx version 4.2 (Waters).

### Liquid chromatography coupled to mass spectrometry

Reversed-phase chromatography was performed using an Agilent 1290 Infinity UHPLC system (Agilent Technologies) coupled to a quadrupole-time-of-flight mass spectrometer (6530 Q-ToF, Agilent Technologies). AncAmor Exo9 and AncOpis Exo9 were diluted to 0.068 mg/mL and 0.056 mg/mL, respectively, in buffer A (0.1% (v/v) formic acid in LC-MS grade water), and 20 µL sample was injected onto a 2.1 mm x 12.5 mm Zorbax 5 µm 300SB-C3 guard column, with column oven temperature maintained at 40 °C. Chromatography began at 90% buffer A, 10% buffer B (0.1% (v/v) formic acid in acetonitrile) at a flow rate of 1.0 mL/min for 3 s. A linear gradient was performed from 10% to 80% buffer B over 17.4 s, followed by a second linear gradient from 80% to 95% buffer B over 1.2 s. Elution was performed isocratically at 95% buffer B over 23.4 s, followed by re-equilibration to the initial conditions over 1.2 s and an isocratic hold for 43.8 s. The mass spectrometer was configured using the standard electrospray ionisation source and operated in positive ion mode. The capillary source used the following parameters: capillary potential 4000 V, nebulizer pressure 60 psi, source temperature of 350 °C, and drying gas flow rate 12 L/min. Ion optic potentials were: skimmer 65 V, fragmentor 250 V, octopole radio frequency 750 V. Data was analysed using Agilent MassHunter Qualitative Analysis B.07.00 software (Agilent Technologies Inc.).

### RNA oligonucleotide preparation

HPLC-grade RNA oligonucleotides were ordered from Eurofins Genomics or Sigma-Aldrich, dissolved in RNase-free water (Takara Bio) to a stock concentration of 100 µM, aliquoted, and stored at −20 °C until usage. RNA probes containing double-stranded regions were prepared by mixing oligos complementary sequences at a concentration of 40 µM in hybridization buffer (10 mM Tris-HCl, pH 8.0, 20 mM KCl). The probes were annealed in a PCR cycler by heating at 95 °C for 7 min and gradually cooling down to 21 °C over 55 min (1.5 °C reduction steps, holding temperature for 1 min per step). Once annealed, the probes were diluted to working concentrations using RNase-free water or Milli-Q water and stored at room temperature until usage. RNAs were freshly annealed from frozen stocks before each experiment. Single-stranded (ss) RNAs that were used to compare ssRNA and dsRNA were prepared similarly, while omitting the addition of complementary oligonucleotides. RNA samples for MP analysis were diluted in SEC100 buffer instead of water. Oligonucleotide sequences are provided in Supplementary Table S4.

### Ribonuclease assays

Ribonuclease (RNase) assays were performed in reaction buffer (RB) containing 20 mM Tris-Cl pH 8.0, 50 mM KCl, 5 mM MgCl_2_, 1 mM DTT, 10% glycerol, 10 mM NaH_2_PO_4_ (pH 7.8). For initial RNase assays (Fig. 2), a concentration of 10 MgCl_2_ was used, which was later replaced with 5 MgCl_2_ because it improved RNase activity of Rrp44. Proteins were diluted to working concentrations using SEC100. To initiate RNA degradation, FAM-labeled RNA was diluted in RB to 10 nM, Exo9 or Rrp44 was added to a concentration of 250 nM, and the reaction was incubated at 37 °C. At indicated time points, 6 µL of the reaction was removed from the tube and stopped by adding 6 µL of formamide loading buffer containing 98% formamide, 10 mM EDTA, 0.1% bromophenol blue, 0.1% xylene cyanol. For control reactions, an equal volume (typically 1 µL per 10 µL total reaction) of SEC100 buffer was added instead of RNase, and the reaction was incubated at 37 °C until the last time point of the experiment was taken. For control reactions without phosphate, an equal amount of water was added instead of NaH_2_PO_4_. For experiments using monomeric or heterodimeric Exo9 subunits, the final protein concentration in the reaction mix was 1 µM. Proteins were pre-diluted to 60 µM in SEC350 buffer. The samples were diluted further to 10 µM using SEC350, 1 µL was added to 9 µL of reaction mix, containing 10 nM of RNA, and incubated at 37 °C for 30 min. Exo9 activity of these samples was tested by mixing equal volumes (1 µL each) of proteins (60 µM), corresponding to 10 µM of Exo9 in the pre-mix, and 1 µM in the reaction mix.

Reactions were analyzed via denaturing PAGE. Denaturing gels (15% in 1x TBE and 8.3 M urea) were hand-casted using a DNA gel sequencing buffer set (ROTIPHORESE, Carl Roth) and the Mini-PROTEAN Tetra (Bio-Rad) system, which was also used for PAGE. For experiments using ssRNA, PAGE was performed at 150 V using 0.5x TBE. Gels were pre-run in 0.5x TBE for 30 min, 10 µL of sample (mixed with loading buffer) was loaded, and the gels were run at 150 V for 60 min. For experiments using dsRNA, the samples were heated at 95 °C for 3 min, and PAGE was performed using 1x TBE at 50 °C. The gels were imaged using a ChemiDoc MP imaging system (Bio-Rad) and the fluorescent signal of the FAM-labeled RNA was detected for 45 s using the Alexa Fluor 488 channel. All RNase experiments were independently repeated at least twice.

For analysis and quantification of RNase assays, RNA band intensities were quantified via Image Lab (Bio-Rad) and plotted using GraphPad Prism. To plot relative RNA intensities, the fluorescent signal of the full-length RNA in each lane was divided by the sum of full-length and degraded RNA (final product, lowest band). RNA decay was fitted using the one-phase decay model in GraphPad. All experiments used for quantification were independently repeated three times.

### Electrophoretic mobility shift assays

Electrophoretic mobility shift assays (EMSA) were performed using native PAGE. Proteins (250 nM) were incubated with FAM-labeled RNA (10 nM) on ice or at 37 °C for at least 30 minutes in binding buffer (20 mM Tris-Cl, pH 8.0, 50 mM KCl, 5 mM MgCl_2_, 1 mM DTT, 10% glycerol), and 9 µL were loaded onto 4% native TBE gels, pre-run at 80 V in 0.5 x TBE at 4 °C for 30 min. For EMSAs used to monitor RNA binding of reconstituted Exo10 complexes, samples were directly taken from the SEC fractions, glycerol was added to a concentration of 10%, and 5 µL were loaded on the native gels. Gels were run at 80 V in 0.5 x TBE at 4 °C for 80 min and imaged as described under ribonuclease assays.

### Cryo-EM sample preparation

AncAmor Exo9 at a concentration of 1.5 µM was incubated with an equimolar amount of dsRNA containing RNA (oligonucleotide 1: 5’-[FAM] AUU CUA UCC CAG CGU CGU AUC UAU CCA AAA UUA AAC AAU AAU CAA UUA CAG UCC CUU UA-3’, oligonucleotide 2: 5’-GAU ACG ACG CUG GGA UAG-3’), annealed as described under ‘RNA oligonucleotide preparation’ and diluted in SEC100 to working concentration. Exo9 and RNA were incubated for 90 min on ice, and four buffer conditions were used for sample preparation. For condition A, Exo9 and RNA were incubated in SEC100 buffer. For condition B, Exo9 and RNA were incubated in buffer containing 20 mM Tris-Cl, pH 8.0,150 mM NaCl, 2 mM MgCl_2_, 0.8 mM DTT. For condition C, Exo9 and RNA were incubated in buffer containing 20 mM Tris-Cl pH 8.0, 80 mM NaCl, 50 mM KCl, 10 mM MgCl_2_, 0.8 mM DTT. For condition D, Exo9 at a concentration of 5.7 µM was incubated with an equimolar amount of RNA in buffer containing 18 mM Tris-Cl pH 8.0,118 mM NaCl, 0.7 mM DTT, 3.8 mM CHAPSO. CHAPSO was dissolved in water to a stock concentration of 40 mM and freshly added before plunge-freezing. AncOpis Exo9 sample was prepared like AncAmor Exo9 under condition C (1.5 µM of Exo9 and RNA, 90 min incubation on ice, buffer containing 20 mM Tris-Cl pH 8.0, 80 mM NaCl, 50 mM KCl, 10 mM MgCl_2_, 0.8 mM DTT). For AncAmorExo10, the frozen complex, prepared as described under ‘RNA exosome reconstitution’, was thawed, diluted to a concentration of 1.5 µM with SEC100 buffer, and directly used for cryo-EM grid preparation.

Cryo-EM samples were prepared by applying 3 µL of sample to C-Flat 1.2/1.3-3Cu-50 grids (Electron Microscopy Sciences), freshly glow-discharged via a PELCO easiGlow system (Ted Pella, Inc.). The grids were plunge-frozen using a Vitrobot Mark IV (Thermo Fisher Scientific), set to 100% humidity, 4 °C. For AncAmor and AncOpis Exo9, the following blotting parameters were used: wait time 0 s, blot force 20 and blot time 4 s. For AncAmor Exo10, the following blotting parameters were used: wait time 0 s, blot force −2 and blot time 8.5 s. Cryo-EM AutoGrids (NanoSoft) were prepared using an auto grid assembly workstation (Thermo Fisher Scientific).

### Cryo-EM data collection and processing

Cryo-EM data were recorded on a Titan Krios G3i transmission electron microscope (Thermo Scientific), operated at 300 keV, and equipped with a BioQuantum imaging filter (Gatan) and K3 direct electron detector (Gatan). Supplementary Table S3 lists cryo-EM data collection parameters. Automated data collection was performed using EPU. For AncAmor Exo9, four datasets (conditions A-D, one dataset per condition) were collected and processed. Because all four datasets, when processed separately, yielded high-resolution cryo-EM reconstructions that didn’t show any obvious differences when we compared them, the high-quality particles from each dataset were combined and processed further. For AncOpis Exo9, one dataset was collected and processed. For AncAmor Exo10, one was collected and processed. Cryo-EM data were processed with CryoSPARC^45^. The cryo-EM processing strategies are outlined in Extended Data Figure 5 (AncAmor Exo9); Extended Data Figure 6 (AncOpis Exo9), and Extended Data Figure 10 (AncAmor Exo10). In brief, movies were motion corrected using Patch Motion Correction, and the results were saved in 16-bit floating point. Contrast transfer function (CTF)-correction was performed using Patch CTF Estimation. Particles were initially picked using Blob Picker and subjected to 2D classification. Selected 2D classes were used for Template Picker and, following iterative rounds of 2D classification, used for particle picking via Topaz^65^. Initial volumes were generated using Ab-Initio Reconstruction with multiple classes and used for Heterogeneous Refinement. Selected particles from higher-quality reconstructions were subjected to Non-uniform Refinement^66^ and 3D classification. Classes featuring cryo-EM density for C-terminal Csl4 were selected and processed further. For AncAmor Exo9 datasets, cryo-EM reconstructions with improved densities for RNA were obtained through iterative rounds of 3D classification with masks covering the RNA. The RNA-unbound AncAmor Exo9 reconstruction was obtained through global 3D classification without masks. AncAmor Exo9 and Exo10 were also subjected to 3D Flexible Refinement^67^. AncAmor Exo10 was also subjected to Local Refinement using a mask that covered the C-terminus of Rrp44. Cryo-EM maps were sharpened with EMReady^43^

### Structural model building, refinement, and analysis

Cryo-EM maps were visualized and analyzed with UCSF ChimeraX^68^. Initial structural models of RNA-bound Exo9 and Exo10 complexes were predicted with AlphaFold 3^69^ and fitted into their cryo-EM maps in ChimeraX. Coot^70,71^ was used for structural model building. First, the predicted models of the individual polypeptides were real-space refined with applied distance restraints, obtained through ProSMART^72^ and implemented in Coot. The map-model fit was inspected and adjusted via real-space refinement with Ramachandran^73^ restraints. To analyze the conformational transition between RNA-bound and unbound AncAmor Exo9, the structural model of RNA-bound AncAmor Exo9 was fitted to the RNA-unbound cryo-EM with Namdinator^74^. All models were iteratively refined via Phenix^75^. Structural model and refinement statistics, derived by MolProbity^76^ (implemented in Phenix), are provided in Supplementary Table S3.

## Data availability

Cryo-EM maps will be deposited at the Electron Microscopy Data Bank (EMDB). Atomic models of human Pol III have been deposited at the Protein Data Bank.

## Acknowledgements

We thank the Central Electron Microscopy Facility at the Max Planck Institute of Biophysics for their expertise and instrument access. We thank C. Thölken, R. Sitt, and M. Lechner for assistance with data transfer and software installation, and for setting up and maintaining the high-performance computing environment. We thank S. Garg for computational assistance to streamline ASR. M.G. and G.K.A.H. were supported by the Max Planck Society. M.G. acknowledges support by the Peter und Traudl Engelhorn Stiftung and the Microcosm Earth Center. F.D.N.A was supported by funding from the Biotechnology and Biological Sciences Research Council (UKRI-BBSRC) [grant number BB/T008784/1]. G.K.A.H. acknowledges funding from a LOEWE Excellence Professorship. J.M.S. acknowledges funding from the Deutsche Forschungsgemeinschaft for an Emmy Noether grant (SCHU 3364/1-1). This work is supported by ERC grant (EVOCATION, 101040472) awarded to G.K.A.H. This research was funded by the European Union. Views and opinions expressed are however those of the author(s) only and do not necessarily reflect those of the European Union or the European Research Council Executive Agency. Neither the European Union nor the granting authority can be held responsible for them.

## Contributions

M.G. and G.K.A.H. conceived this project and planned experiments. M.G. performed phylogenetic analysis, ancestral sequence reconstruction, molecular cloning, protein purification and complex reconstitution, MP analysis, RNA degradation experiments, cryo-EM data processing, structural model building, refinement, and interpreted the data. F.D.N.A. performed native MS and LC-MS experiments. S.P. prepared cryo-EM grids and collected cryo-EM data. L.A. assisted with molecular cloning and protein purification and performed RNA degradation experiments. J.M.S. contributed to project conceptualization. F.D.N.A. and J.L.P.B. planned and analyzed MS experiments. G.K.A.H. supervised this project and interpreted the data. M.G. and G.K.A.H. wrote the manuscript with input from the other authors.

---

**Extended Data Figure 1.**
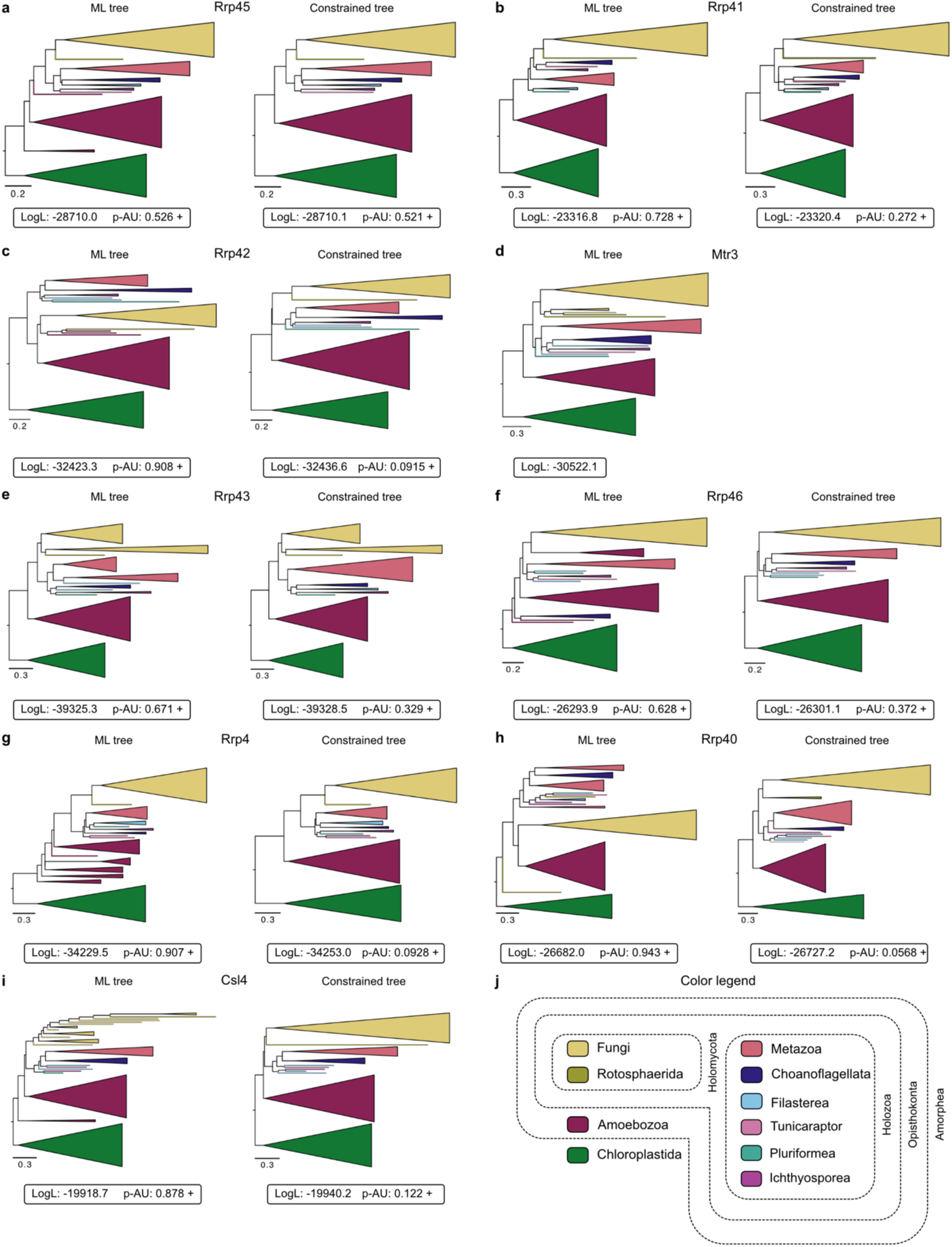
Phylogenies of RNA exosome core subunits. **a–i,** Phylogenies of Exo9 subunits. Shown are the maximum likelihood (ML) trees (left) and the constrained trees (right), together with Log-likelihood (LogL) values and p-values of the approximately unbiased (AU) tests^27^ (p-AU). The plus signs mark 95% confidence sets, indicating that the constrained trees are not rejected by the AU tests. Scale bars: average substitutions per site. **j,** Color legend for taxonomic coloring of the clades or branches and grouping of clades.

**Extended Data Figure 2.**
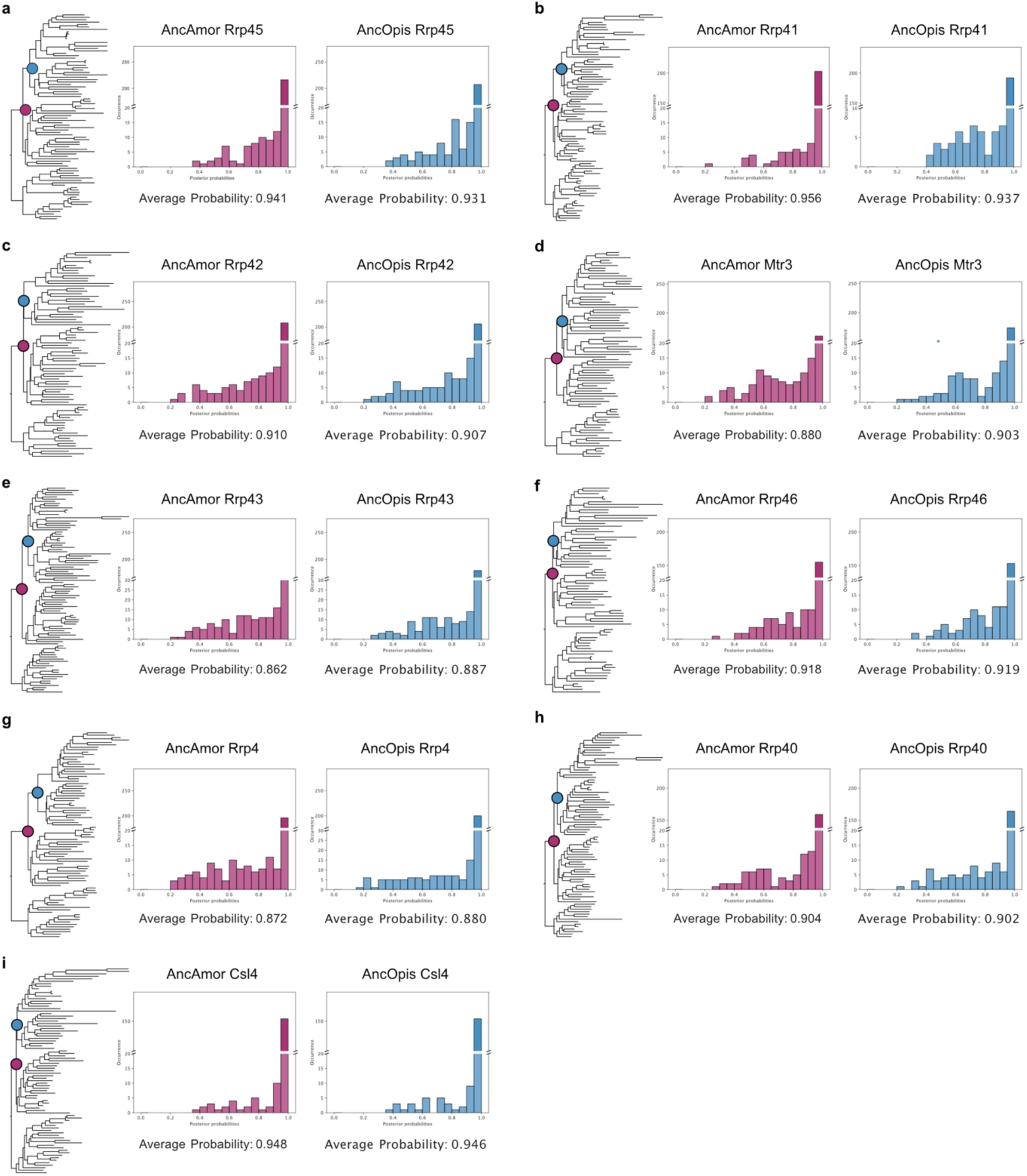
Ancestral sequence reconstruction (ASR) of ancestral Exo9 proteins. **a–i,** ASR of individual Exo9 subunits. Resurrected nodes (AncAmor and AncOpis) are labeled in the phylogenetic trees (left). Shown are posterior probability distributions of AncAmor (middle) and AncOpis (right) reconstructions. Posterior probabilities correspond to the likelihood of ancestral states at each site and provide a measure of confidence in the reconstructed states. The x-axis shows the range of posterior probabilities. The y-axis shows the occurrence of sites with corresponding posterior probabilities. The average posterior probability value across all sites, given below the distribution plots, reflects the overall confidence in the reconstructed sequences.

**Extended Data Figure 3.**
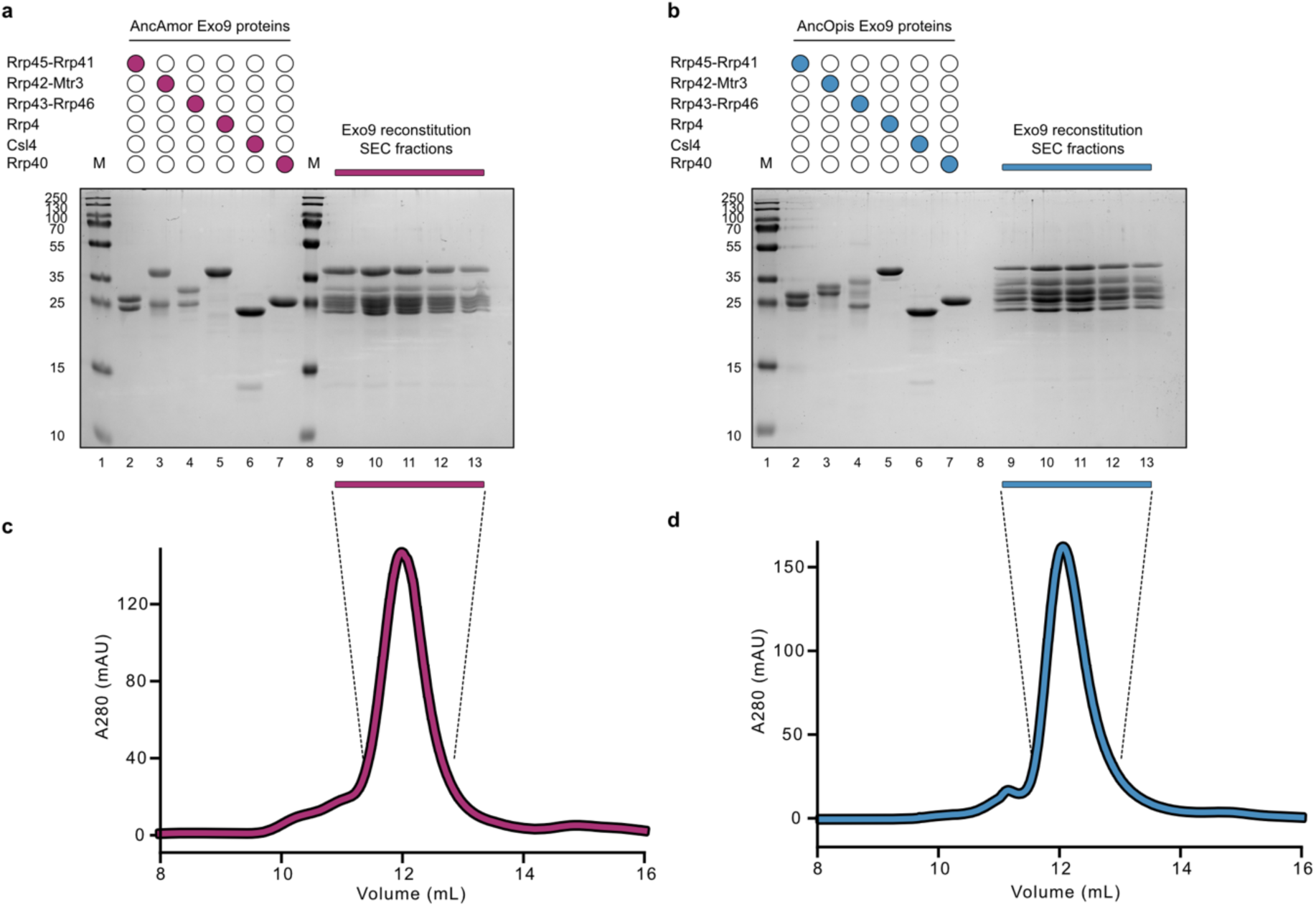
Purification of Exo9 proteins and reconstitution of ancestral Exo9 complexes. **a–b,** Protein gels (12% SDS-PAGE) of purified AncAmor (**a**) and AncOpis (**b**) Exo9 proteins. Dimeric ring proteins (lanes 2–4) and monomeric cap proteins (lanes 5–7) are loaded as indicated above by colored circles. **c–d,** Size-exclusion chromatography (SEC) profiles of reconstituted AncAmor (**c**) and AncOpis (**d**) Exo9 complexes loaded onto an ENrich SEC 650 column. Highlighted peak fractions were loaded on the protein gels shown in (**a, b**) as indicated (lanes 9–13), indicating that the loaded input Exo9 proteins co-migrate as a higher molecular-weight complex over the SEC column.

**Extended Data Figure 4.**
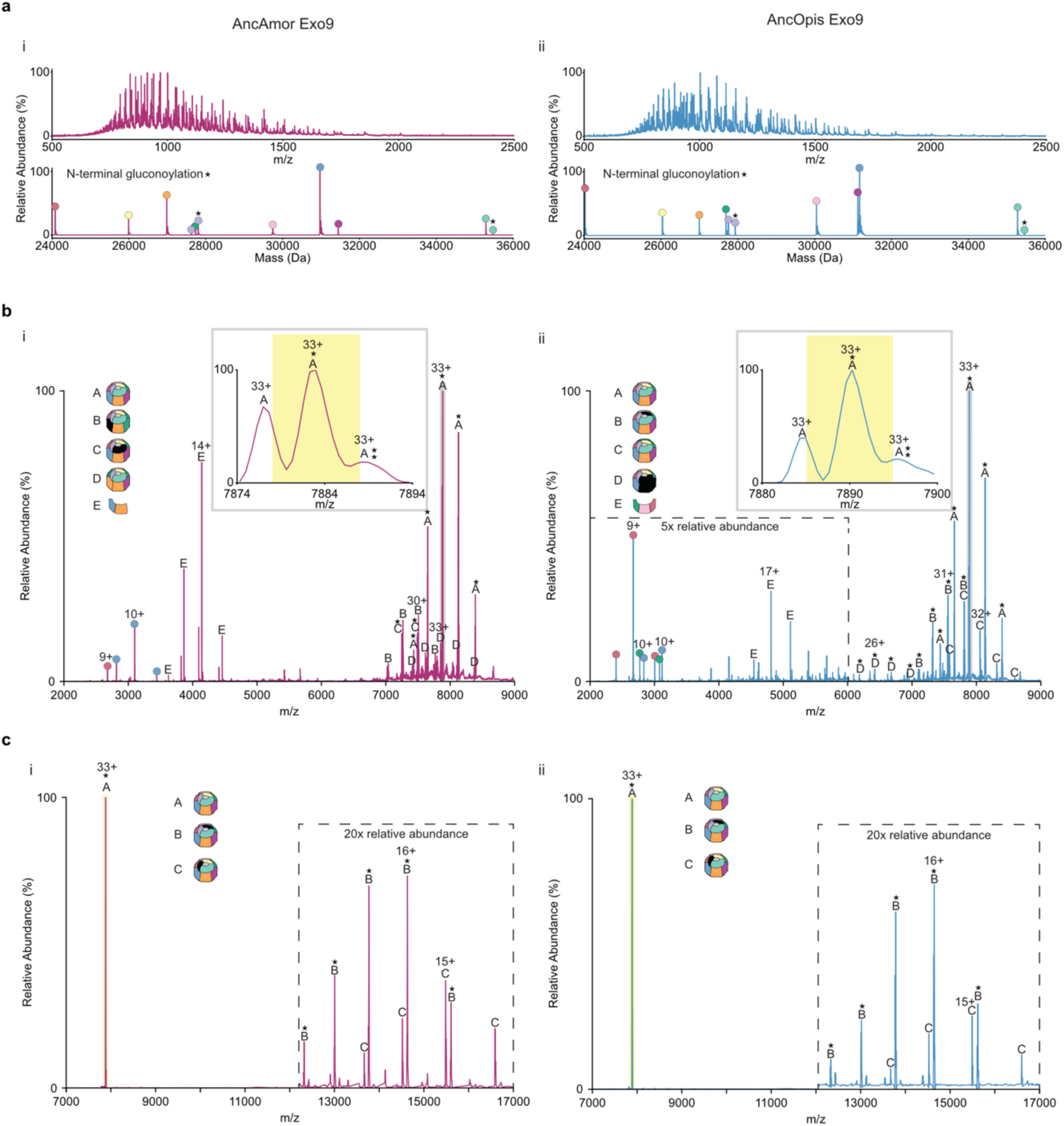
Liquid chromatography-mass spectrometry (LC-MS), native MS, and tandem MS of AncAmor Exo9 and AncOpis Exo9. **a,** LC-MS of AncAmor 9-9 (i) and AncOpis 9-9 (ii) revealed the exact masses of each of the Exo9 subunits, as shown in the deconvolved mass spectra (bottom). Subunits Rrp4 and Rrp40 have the N-terminus Gly-Ser-Ser-[His]_6_- that can be gluconoylated (*) during *E. coli* expression, resulting in a 178 Da mass addition^44^. **b,** Native MS of AncAmor 9-9 (i) and AncOpis 9-9 (ii) revealed that the most abundant complex (species A) has a 1:1:1:1:1:1:1:1:1 stoichiometry. Small fractions of mis-assembled complexes were detected in each sample. Focusing on the 33+ charge state of the correctly assembled nonamer (inset) shows its detection with zero, one (*), or two (**) gluconoylations, with a distribution arising from the gluconoylated fractions of Rrp4 and Rrp40. An ensemble of the singly-gluconoylated nonameric ions within a ±5 *m/z* window (yellow) was selected for tandem MS. **c,** This revealed loss of the CsI4 or Rrp40 subunits and confirmed the 1:1:1:1:1:1:1:1:1 stoichiometry of the nonamer in both AncAmor 9-9 (i) and AncOpis 9-9 (ii).

**Extended Data Figure 5.**
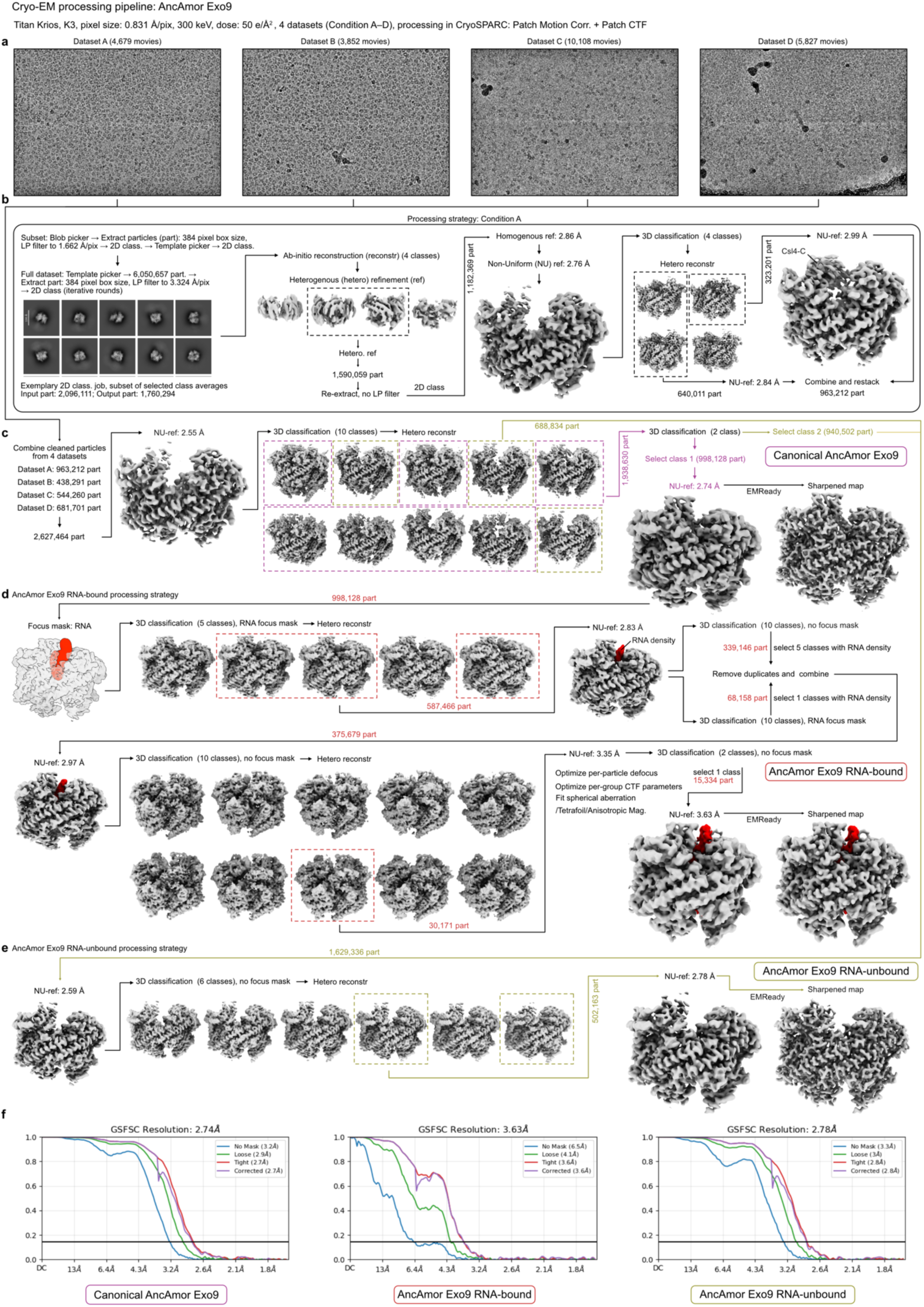
Cryo-EM data processing strategy of AncAmor Exo9 complexes. Collection parameters are provided at the top. Processing was performed via cryoSPARC^45^. Four datasets were recorded. All samples contained similar protein and RNA content, but buffer conditions were different. Dataset A: 20 mM Tris-HCl, pH 8.0, 100 mM NaCl, 1 mM DTT. Dataset B: 20 mM Tris-Cl pH 8.0,150 mM NaCl, 2 mM MgCl_2_, 0.8 mM DTT. Dataset C: 20 mM Tris-Cl pH 8.0, 80 mM NaCl, 50 mM KCl, 10 mM MgCl_2_, 0.8 mM DTT. Dataset D: 18 mM Tris-Cl pH 8.0,118 mM NaCl, 0.7 mM DTT, 3.8 mM CHAPSO. Exo9/RNA concentrations datasets A–C: 1.5 µM; dataset D: 5.7 µM. Shown are the unsharpened cryo-EM maps, except for the EMready^43^-sharpened maps. The number of particles (part) used for processing jobs is indicated. **a,** Representative micrographs (5 Å low-pass filtered) of the collected datasets. The number of collected movies are provided above. **b,** Processing strategy for dataset A. The other datasets were processed similarly. c, Strategy for the canonical AncAmor Exo9 map, derived by combining cleaned particles of the four datasets. Particles used in the canonical map processing are colored purple. d, Strategy of the AncAmor Exo9 RNA-bound map. Particles used for RNA-bound Exo9 processing are colored red. The RNA density was improved through iterative rounds of 3D classification with and without a mask applied to the RNA-containing region. An exemplary mask, colored in red, is shown on the top left. e, Strategy of the AncAmor Exo9 RNA-unbound map. Particles used for RNA-bound Exo9 processing are colored olive. f, Fourier shell correlation (FSC) curves of AncAmor Exo9 canonical (left), AncAmor Exo9 RNA-bound (middle), and AncAmor Exo9 RNA-unbound (right).

**Extended Data Figure 6.**
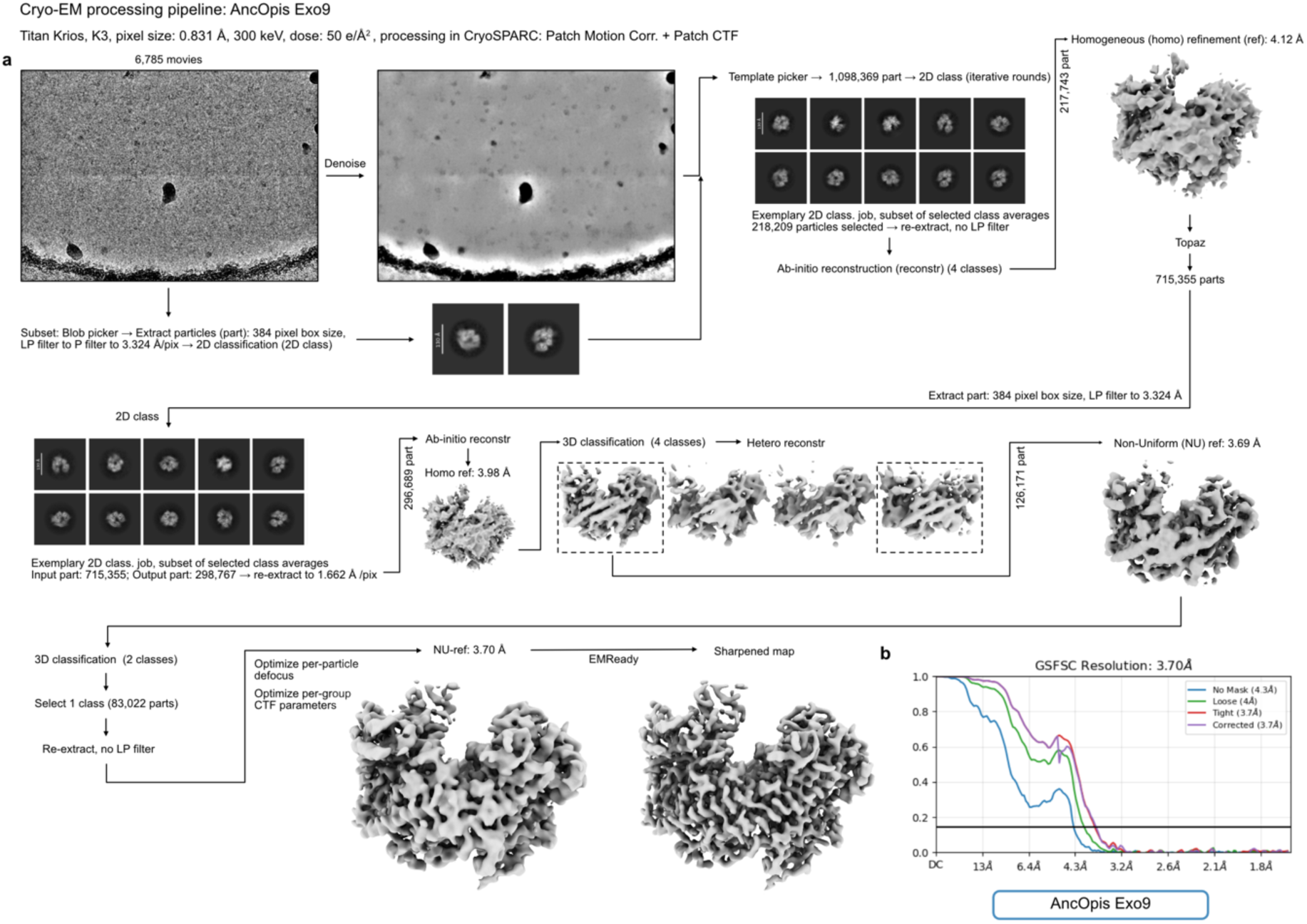
Cryo-EM data processing strategy of the AncOpis Exo9 complex. Collection parameters are provided at the top. Processing was performed via cryoSPARC^45^. Shown are the unsharpened cryo-EM maps, except for the EMready^43^-sharpened maps. **a,** Processing strategy for AncOpis Exo9. The number of collected movies is provided above a representative micrograph, shown with a 5 Å low-pass filter (left) and denoised (right). and of the collected datasets. The number of particles (part) used for processing jobs is indicated. **b,** Fourier shell correlation (FSC) curve of AncOpisExo9.

**Extended Data Figure 7.**
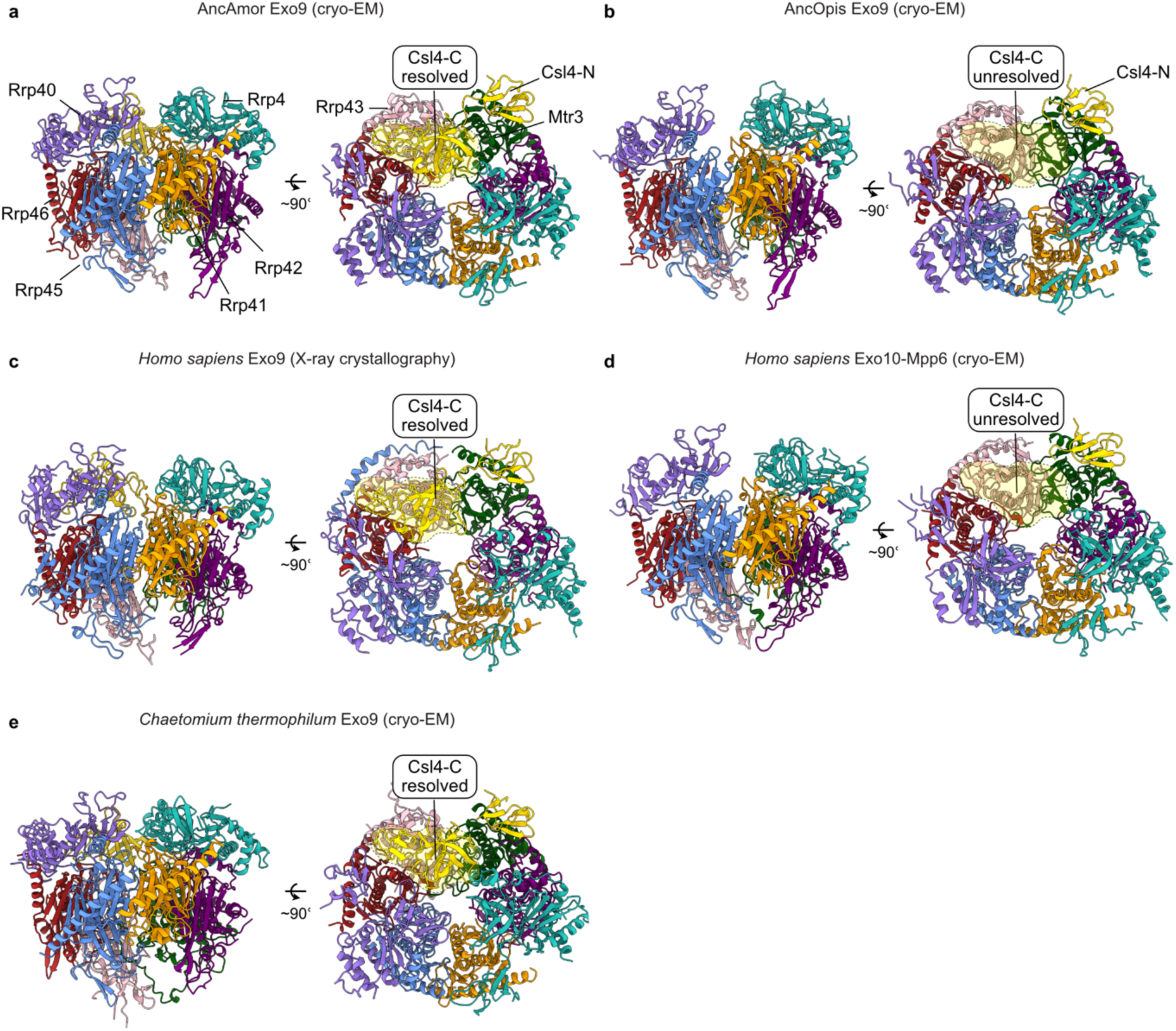
Comparisons between ancestral and selected extant RNA exosome structures. **a–e,** Side and top views of superimposed structural models of ancestral and selected extant RNA exosomes. The ancestral Exo9 structures overall match their extant counterparts, indicating the integrity of resurrected ancestral RNA exosomes. **a,** AncAmor Exo9 (this study). **b,** AncOpis Exo9 (this study). **c,** *Homo sapiens* Exo9 (PDB: 2NN6, Ref^5^). **d,** *H. sapiens* Exo10-Mpp6 (PDB: 6H25, Ref^30^). Rrp44, Mpp6, and RNA chains are hidden for better comparison between Exo9 models. **e,** *Chaetomium thermophilum* Exo9 (PDB: 8R1O, Ref^29^). The Exo9 proteins are labeled in AncAmor Exo9, and the color code is similar for all structures. The C-terminal domain of Csl4 (Csl4-C) is resolved in AncAmor Exo9 (**a**), the X-ray crystallography structure of *H. sapiens* Exo9 (**c**), and *C. thermophilum* Exo9 (**e**). The Csl4-C is unresolved in AncOpis Exo9 (**b**) and H. sapiens Exo10-Mpp6 (**d**). The yellow-colored, transparent shape marks the presence or absence of this domain.

**Extended Data Figure 8.**
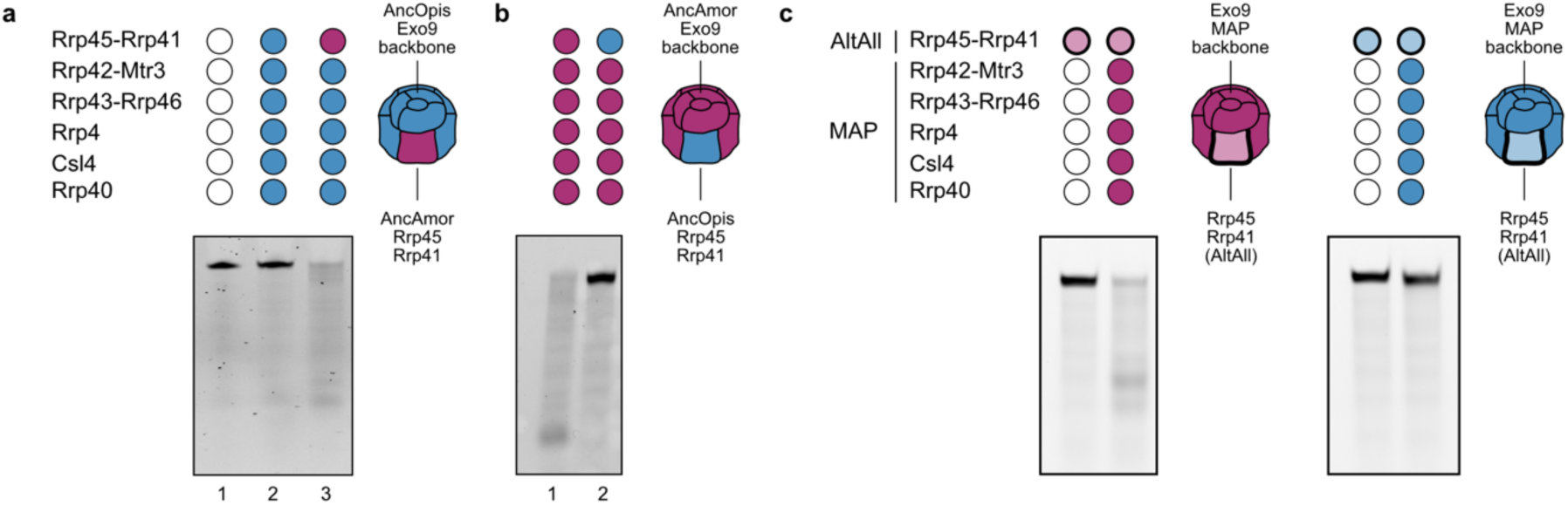
RNase activities of Exo9 with interchanged or alternative Rrp45-Rrp41 ancestors. **a,** RNase assay of AncOpis Exo9, assembled through mixing equimolar amounts of AncOpis Exo9 proteins. AncOpis Rp45-41 was replaced with AncAmor Rrp45-41 in lane 3 as illustrated by the schematic on the right. **b,** RNase assay of AncOpis Exo9, assembled through mixing equimolar amounts of AncAmor Exo9 proteins. AncAmor Rp45-41 was replaced with AncOpis Rrp45-41 (lane 2). **c,** Probing the robustness of ancestral reconstructions. Exo9 complexes were assembled by mixing equimolar amounts of AncAmor (left gel) or AncOpis (right gel) Exo9 proteins. The maximum a posteriori (MAP) ancestors were used for the Exo9 backbones. The alternative, less-likely reconstructions (AltAll) of Rrp45-41 were used, which conferred RNase activity in AncAmor Exo9 and no RNase activity in AncOpis Exo9. The RNA substrate carries a fluorescent carboxyfluorescein (FAM) label on the 5’-end. Gel: denaturing acrylamide gel (15% urea in 1xTBE).

**Extended Data Figure 9.**
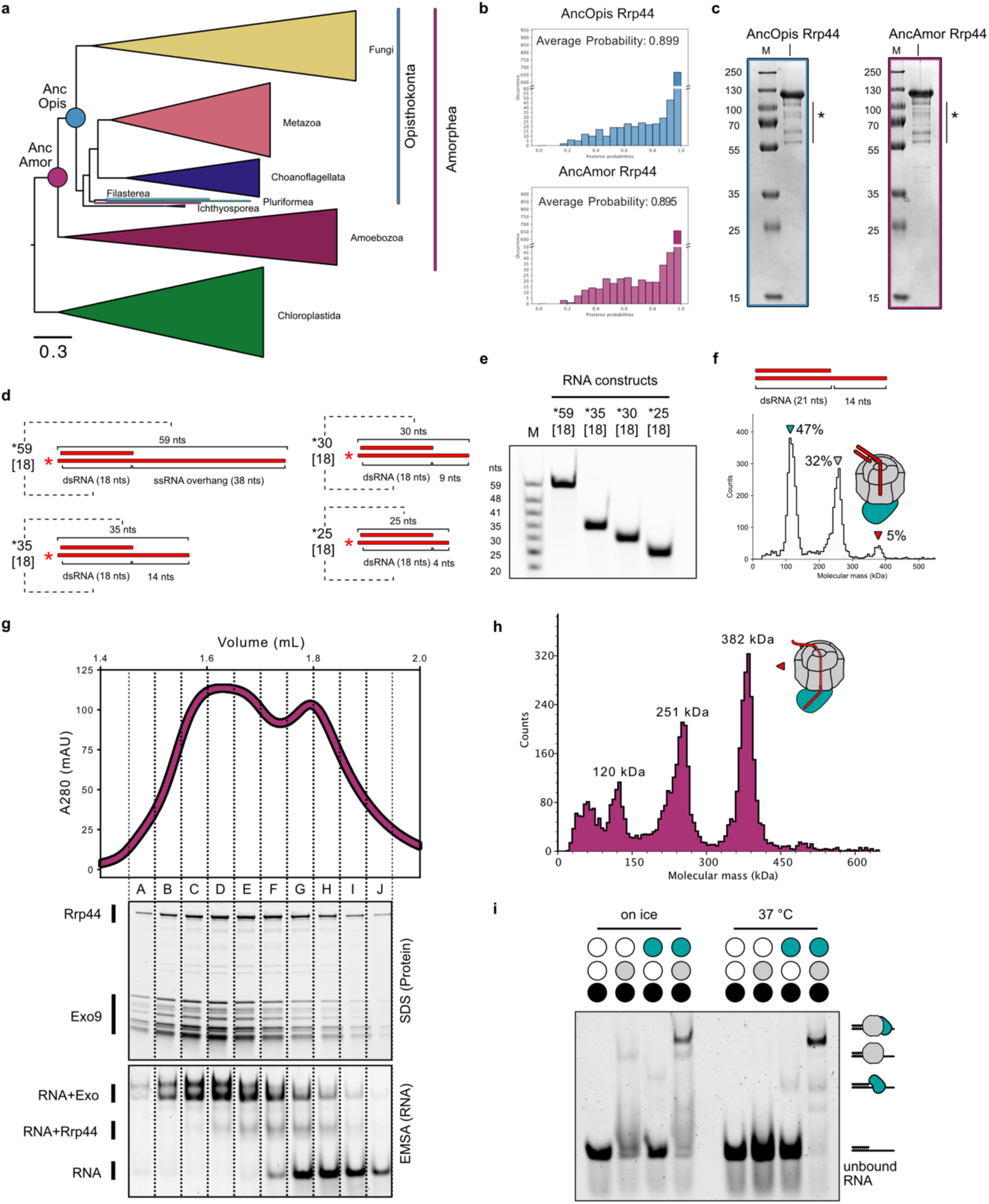
Rrp44 resurrection, interaction with Exo9 and Exo10 reconstitution. **a,** ML tree of Rrp44. Resurrected key nodes are shown as colored circles. Scale bar: average substitutions per site. **b,** Posterior probability distributions of AncOpis (top) and AncAmor (bottom) Rrp44 reconstructions. **c,** Protein gels (12% SDS-PAGE) of purified AncOpis (left) and AncAmor (right) Rrp44. Asterisks mark proteolytic degradation products co-eluting during size-exclusion chromatography (SEC). **d,** Schematic of RNA constructs used for MP analysis. **e,** Denaturing RNA gel of RNA constructs shown in (**d**). **f,** Mass photometry (MP) analysis of AncAmor Exo10, formed on an alternative RNA construct lacking a fluorescent tag and a longer hybridization oligo. g, Reconstitution of AncAmor Exo10 for cryo-EM. Top: SEC profile. SEC Fractions are labeled with capitalized letters and separated by dashed lines. Middle: EZFluor-stained protein gel of SEC fractions. Bottom: Electrophoretic mobility shift assay (EMSA) of SEC fraction, showing the AmoExo10 complex is fully bound to FAM-labeled RNA. h, MP analysis of reconstituted AncAmor Exo10. i, EMSA of AncAmor Exo9 and Rrp44 on RNA constructs used to probe dsRNA degradation. Incubation on ice (left) or 37 °C (right).

**Extended Data Figure 10.**
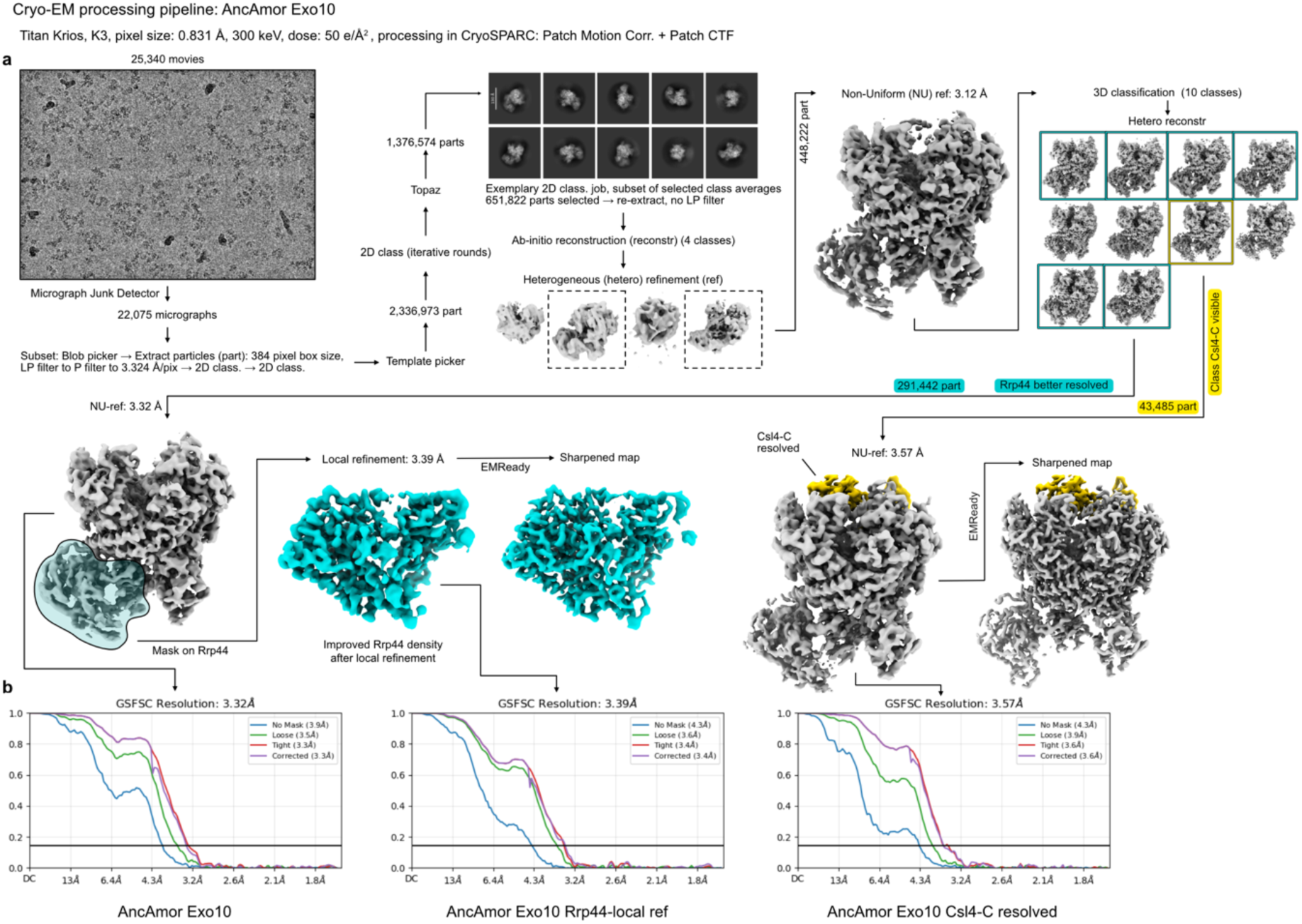
Cryo-EM data processing strategy of the AncAmor Exo10 complex. Collection parameters are provided at the top. Processing was performed via cryoSPARC^45^. Shown are the unsharpened cryo-EM maps, except for the EMready^43^-sharpened maps. **a,** Processing strategy for AncAmor Exo10. The number of collected movies is provided above a representative micrograph (5 Å low-pass filtered). The number of particles (part) used for processing jobs is indicated. Rrp44 density was improved via local refinement with a mask applied to Rrp44. The occupancy of the C-terminal domain of Csl4 (Csl4-C) was improved via 3D classification, as indicated. **b,** Fourier shell correlation (FSC) curves of AncAmor Exo10 reconstructions.

## References

1. Mitchell, P., Petfalski, E., Shevchenko, A., Mann, M. & Tollervey, D. The Exosome: A Conserved Eukaryotic RNA Processing Complex Containing Multiple 3′→5′ Exoribonucleases. Cell 91, 457–466 (1997).

2. Mitchell, P. & Tollervey, D. An NMD Pathway in Yeast Involving Accelerated Deadenylation and Exosome-Mediated 3′→5′ Degradation. Mol. Cell 11, 1405–1413 (2003).

3. Houseley, J. & Tollervey, D. The Many Pathways of RNA Degradation. Cell 136, 763– 776 (2009).

4. Kilchert, C., Wittmann, S. & Vasiljeva, L. The regulation and functions of the nuclear RNA exosome complex. Nat. Rev. Mol. Cell Biol. 17, 227–239 (2016).

5. Liu, Q., Greimann, J. C. & Lima, C. D. Reconstitution, Activities, and Structure of the Eukaryotic RNA Exosome. Cell 127, 1223–1237 (2006).

6. Liu, Q., Greimann, J. C. & Lima, C. D. Reconstitution, Activities, and Structure of the Eukaryotic RNA Exosome. Cell 131, 188–189 (2007).

7. Bonneau, F., Basquin, J., Ebert, J., Lorentzen, E. & Conti, E. The Yeast Exosome Functions as a Macromolecular Cage to Channel RNA Substrates for Degradation. Cell 139, 547–559 (2009).

8. Dziembowski, A., Lorentzen, E., Conti, E. & Séraphin, B. A single subunit, Dis3, is essentially responsible for yeast exosome core activity. Nat. Struct. Mol. Biol. 14, 15– 22 (2007).

9. Briggs, M. W., Burkard, K. T. D. & Butler, J. S. Rrp6p, the Yeast Homologue of the Human PM-Scl 100-kDa Autoantigen, Is Essential for Efficient 5.8 S rRNA 3′ End Formation. Journal of Biological Chemistry 273, 13255–13263 (1998).

10. Wasmuth, E. V. & Lima, C. D. Exo- and Endoribonucleolytic Activities of Yeast Cytoplasmic and Nuclear RNA Exosomes Are Dependent on the Noncatalytic Core and Central Channel. Mol. Cell 48, 133–144 (2012).

11. Makino, D. L., Baumgärtner, M. & Conti, E. Crystal structure of an rna-bound 11-subunit eukaryotic exosome complex. Nature 495, 70–75 (2013).

12. Makino, D. L. et al. RNA degradation paths in a 12-subunit nuclear exosome complex. Nature 524, 54–58 (2015).

13. Zinder, J. C., Wasmuth, E. V. & Lima, C. D. Nuclear RNA Exosome at 3.1 Å Reveals Substrate Specificities, RNA Paths, and Allosteric Inhibition of Rrp44/Dis3. Mol. Cell 64, 734–745 (2016).

14. Burkard, K. T. D. & Butler, J. S. A Nuclear 3′-5′ Exonuclease Involved in mRNA Degradation Interacts with Poly(A) Polymerase and the hnRNA Protein Npl3p. Mol. Cell. Biol. 20, 604–616 (2000).

15. Lebreton, A., Tomecki, R., Dziembowski, A. & Séraphin, B. Endonucleolytic RNA cleavage by a eukaryotic exosome. Nature 456, 993–996 (2008).

16. Evguenieva-Hackenberg, E., Walter, P., Hochleitner, E., Lottspeich, F. & Klug, G. An exosome-like complex in *Sulfolobus solfataricus*. EMBO Rep. 4, 889–893 (2003).

17. Lorentzen, E. et al. The archaeal exosome core is a hexameric ring structure with three catalytic subunits. Nat. Struct. Mol. Biol. 12, 575–581 (2005).

18. Lorentzen, E. & Conti, E. Structural Basis of 3′ End RNA Recognition and Exoribonucleolytic Cleavage by an Exosome RNase PH Core. Mol. Cell 20, 473–481 (2005).

19. Büttner, K., Wenig, K. & Hopfner, K.-P. Structural Framework for the Mechanism of Archaeal Exosomes in RNA Processing. Mol. Cell 20, 461–471 (2005).

20. Walter, P. et al. Characterization of native and reconstituted exosome complexes from the hyperthermophilic archaeon *Sulfolobus solfataricus*. Mol. Microbiol. 62, 1076– 1089 (2006).

21. Audin, M. J. C., Wurm, J. P., Cvetkovic, M. A. & Sprangers, R. The oligomeric architecture of the archaeal exosome is important for processive and efficient RNA degradation. Nucleic Acids Res. 44, 2962–2973 (2016).

22. Sikorska, N., Zuber, H., Gobert, A., Lange, H. & Gagliardi, D. RNA degradation by the plant RNA exosome involves both phosphorolytic and hydrolytic activities. Nat. Commun. 8, 2162 (2017).

23. Chekanova, J. A. et al. Genome-Wide High-Resolution Mapping of Exosome Substrates Reveals Hidden Features in the Arabidopsis Transcriptome. Cell 131, 1340–1353 (2007).

24. Lange, H. et al. The RNA Helicases AtMTR4 and HEN2 Target Specific Subsets of Nuclear Transcripts for Degradation by the Nuclear Exosome in Arabidopsis thaliana. PLoS Genet. 10, e1004564 (2014).

25. Lange, H. et al. RST1 and RIPR connect the cytosolic RNA exosome to the Ski complex in Arabidopsis. Nat. Commun. 10, 3871 (2019).

26. Hochberg, G. K. A. & Thornton, J. W. Reconstructing Ancient Proteins to Understand the Causes of Structure and Function. Annu. Rev. Biophys. 46, 247–269 (2017).

27. Shimodaira, H. An Approximately Unbiased Test of Phylogenetic Tree Selection. Syst. Biol. 51, 492–508 (2002).

28. Greimann, J. C. & Lima, C. D. Reconstitution of RNA exosomes from human and Saccharomyces cerevisiae cloning, expression, purification, and activity assays. Methods Enzymol. 448, 185–210 (2008).

29. Liebau, J. et al. 4D structural biology–quantitative dynamics in the eukaryotic RNA exosome complex. Nat. Commun. 16, 7896 (2025).

30. Gerlach, P. et al. Distinct and evolutionary conserved structural features of the human nuclear exosome complex. Elife 7, (2018).

31. Weick, E.-M. et al. Helicase-Dependent RNA Decay Illuminated by a Cryo-EM Structure of a Human Nuclear RNA Exosome-MTR4 Complex. Cell 173, 1663–1677.e21 (2018).

32. Jacob, F. Evolution and Tinkering. Science (1979). 196, 1161–1166 (1977).

33. Liu, J.-J. et al. Visualization of distinct substrate-recruitment pathways in the yeast exosome by EM. Nat. Struct. Mol. Biol. 21, 95–102 (2014).

34. Liu, J.-J. et al. CryoEM structure of yeast cytoplasmic exosome complex. Cell Res. 26, 822–837 (2016).

35. Wasmuth, E. V, Zinder, J. C., Zattas, D., Das, M. & Lima, C. D. Structure and reconstitution of yeast Mpp6-nuclear exosome complexes reveals that Mpp6 stimulates RNA decay and recruits the Mtr4 helicase. Elife 6, (2017).

36. Kowalinski, E. et al. Structure of a Cytoplasmic 11-Subunit RNA Exosome Complex. Mol. Cell 63, 125–134 (2016).

37. Schuller, J. M., Falk, S., Fromm, L., Hurt, E. & Conti, E. Structure of the nuclear exosome captured on a maturing preribosome. Science (1979). 360, 219–222 (2018).

38. de la Cruz, J. Dob1p (Mtr4p) is a putative ATP-dependent RNA helicase required for the 3’ end formation of 5.8S rRNA in Saccharomyces cerevisiae. EMBO J. 17, 1128– 1140 (1998).

39. LaCava, J. et al. RNA Degradation by the Exosome Is Promoted by a Nuclear Polyadenylation Complex. Cell 121, 713–724 (2005).

40. Wyers, F. et al. Cryptic Pol II Transcripts Are Degraded by a Nuclear Quality Control Pathway Involving a New Poly(A) Polymerase. Cell 121, 725–737 (2005).

41. Lubas, M. et al. Interaction Profiling Identifies the Human Nuclear Exosome Targeting Complex. Mol. Cell 43, 624–637 (2011).

42. Meola, N. et al. Identification of a Nuclear Exosome Decay Pathway for Processed Transcripts. Mol. Cell 64, 520–533 (2016).

43. He, J., Li, T. & Huang, S.-Y. Improvement of cryo-EM maps by simultaneous local and non-local deep learning. Nat. Commun. 14, 3217 (2023).

44. Geoghegan, K. F. et al. Spontaneous α-N-6-Phosphogluconoylation of a “His Tag” inEscherichia coli:The Cause of Extra Mass of 258 or 178 Da in Fusion Proteins. Anal. Biochem. 267, 169–184 (1999).

45. Punjani, A., Rubinstein, J. L., Fleet, D. J. & Brubaker, M. A. cryoSPARC: algorithms for rapid unsupervised cryo-EM structure determination. Nat. Methods 14, 290–296 (2017).

46. Altschul, S. F., Gish, W., Miller, W., Myers, E. W. & Lipman, D. J. Basic local alignment search tool. J. Mol. Biol. 215, 403–10 (1990).

47. Richter, D. J., et al. EukProt: A database of genome-scale predicted proteins across the diversity of eukaryotes. Peer Community Journal 2, (2022).

48. Priyam, A. et al. Sequenceserver: A Modern Graphical User Interface for Custom BLAST Databases. Mol. Biol. Evol. 36, 2922–2924 (2019).

49. Katoh, K. & Standley, D. M. MAFFT multiple sequence alignment software version 7: improvements in performance and usability. Mol. Biol. Evol. 30, 772–80 (2013).

50. Rozewicki, J., Li, S., Amada, K. M., Standley, D. M. & Katoh, K. MAFFT-DASH: integrated protein sequence and structural alignment. Nucleic Acids Res. 47, W5–W10 (2019).

51. Larsson, A. AliView: a fast and lightweight alignment viewer and editor for large datasets. Bioinformatics 30, 3276–8 (2014).

52. Edgar, R. C. MUSCLE: multiple sequence alignment with high accuracy and high throughput. Nucleic Acids Res. 32, 1792–7 (2004).

53. Minh, B. Q. et al. IQ-TREE 2: New Models and Efficient Methods for Phylogenetic Inference in the Genomic Era. Mol. Biol. Evol. 37, 1530–1534 (2020).

54. Kalyaanamoorthy, S., Minh, B. Q., Wong, T. K. F., von Haeseler, A. & Jermiin, L. S. ModelFinder: fast model selection for accurate phylogenetic estimates. Nat. Methods 14, 587–589 (2017).

55. Hoang, D. T., Chernomor, O., von Haeseler, A., Minh, B. Q. & Vinh, L. S. UFBoot2: Improving the Ultrafast Bootstrap Approximation. Mol. Biol. Evol. 35, 518–522 (2018).

56. Guindon, S. et al. New algorithms and methods to estimate maximum-likelihood phylogenies: assessing the performance of PhyML 3.0. Syst. Biol. 59, 307–21 (2010).

57. Rambaut, A. FigTree v.1.4.4. Institute of Evolutionary Biology, University of Edinburgh, Edinburgh (2022).

58. Maddison, W. P. & Maddison, D. R. Mesquite: a modular system for evolutionary analysis. Version 3.7. (2021).

59. Ishikawa, S. A., Zhukova, A., Iwasaki, W. & Gascuel, O. A Fast Likelihood Method to Reconstruct and Visualize Ancestral Scenarios. Mol. Biol. Evol. 36, 2069–2085 (2019).

60. Yang, Z., Kumar, S. & Nei, M. A new method of inference of ancestral nucleotide and amino acid sequences. Genetics 141, 1641–50 (1995).

61. de Vries, S. T., Kley, L. & Schindler, D. Use of a Golden Gate Plasmid Set Enabling Scarless MoClo-Compatible Transcription Unit Assembly. in 105–131 (2025). doi:10.1007/978-1-0716-4220-7_7.

62. Edelheit, O., Hanukoglu, A. & Hanukoglu, I. Simple and efficient site-directed mutagenesis using two single-primer reactions in parallel to generate mutants for protein structure-function studies. BMC Biotechnol. 9, 61 (2009).

63. Kondrat, F. D. L., Struwe, W. B. & Benesch, J. L. P. Native Mass Spectrometry: Towards High-Throughput Structural Proteomics. in 349–371 (2015). doi:10.1007/978-1-4939-2230-7_18.

64. Marty, M. T. et al. Bayesian Deconvolution of Mass and Ion Mobility Spectra: From Binary Interactions to Polydisperse Ensembles. Anal. Chem. 87, 4370–4376 (2015).

65. Bepler, T. et al. Positive-unlabeled convolutional neural networks for particle picking in cryo-electron micrographs. Nat. Methods 16, 1153–1160 (2019).

66. Punjani, A., Zhang, H. & Fleet, D. J. Non-uniform refinement: adaptive regularization improves single-particle cryo-EM reconstruction. Nat. Methods 17, 1214–1221 (2020).

67. Punjani, A. & Fleet, D. J. 3DFlex: determining structure and motion of flexible proteins from cryo-EM. Nat. Methods 20, 860–870 (2023).

68. Goddard, T. D. et al. UCSF ChimeraX: Meeting modern challenges in visualization and analysis. Protein Science 27, 14–25 (2018).

69. Abramson, J. et al. Accurate structure prediction of biomolecular interactions with AlphaFold 3. Nature 630, 493–500 (2024).

70. Emsley, P., Lohkamp, B., Scott, W. G. & Cowtan, K. Features and development of Coot. Acta Crystallogr. D Biol. Crystallogr. 66, 486–501 (2010).

71. Casañal, A., Lohkamp, B. & Emsley, P. Current developments in Coot for macromolecular model building of Electron Cryo-microscopy and Crystallographic Data. Protein Science 29, 1069–1078 (2020).

72. Nicholls, R. A., Long, F. & Murshudov, G. N. Low-resolution refinement tools in REFMAC 5. Acta Crystallogr. D Biol. Crystallogr. 68, 404–417 (2012).

73. Ramachandran, G. N., Ramakrishnan, C. & Sasisekharan, V. Stereochemistry of polypeptide chain configurations. J. Mol. Biol. 7, 95–99 (1963).

74. Kidmose, R. T., et al. *Namdinator* – automatic molecular dynamics flexible fitting of structural models into cryo-EM and crystallography experimental maps. IUCrJ 6, 526– 531 (2019).

75. Liebschner, D. et al. Macromolecular structure determination using X-rays, neutrons and electrons: recent developments in Phenix. Acta Crystallogr. D Struct. Biol. 75, 861–877 (2019).

76. Davis, I. W. et al. MolProbity: all-atom contacts and structure validation for proteins and nucleic acids. Nucleic Acids Res. 35, W375–W383 (2007).

