## Supplementary information for "Coordination of sequential RNase activities in an ancient molecular machine"

**Supplementary Table S1 | List of the AncAmor 9-9 and AncOpis 9-9 subunits and their masses measured using LC-MS and native MS (nMS).** Subunits Rrp4 and Rrp40 have the N-terminus Gly-Ser-Ser-[His]6- that can be gluconoylated (*) during E. coli expression, resulting in a 178 Da mass addition.[1] Subunits that were not detected using nMS are indicated with a dash.

| Subunit | AncAmor Exo9 | | AncOpis Exo9 | |
|---|---|---|---|---|
| | Mass (LC-MS) /Da | Mass (nMS) /Da | Mass (LC-MS) /Da | Mass (nMS) /Da |
| Rrp4 | 35292 | — | 35279 | — |
| Rrp4* | 35471 | — | 35458 | — |
| Rrp40 | 27626 | — | 27755 | — |
| Rrp40* | 27804 | — | 27934 | — |
| Rrp41 | 26985 | — | 26998 | — |
| Rrp42 | 31448 | — | 31123 | — |
| Rrp43 | 29747 | — | 30047 | — |
| Rrp45 | 30973 | 30974 | 31168 | 31168 |
| Rrp46 | 24080 | 24080 | 24023 | 24023 |
| Csl4 | 25992 | — | 26038 | — |
| Mtr3 | 27735 | — | 27693 | 27693 |

**Supplementary Table S2 | List of AncAmor 9-9 and AncOpis 9-9 protein species and their masses and abundances measured using native MS.** Species were detected and mass and abundances were measured using UniDec version 6.0.3.[2] Mass was sampled every 1.0 Da, the peak detection range was set to 50.0 Da, and the peak detection threshold was set to 0.03. Species were assigned using the subunit masses listed in Supplementary Table 1. Any detected species that could not be assigned were ignored. The Mtr3 monomer (green) in AncOpis Exo9 was below the peak detection threshold, so its mass was deconvolved using MassLynx version 4.2 (Waters). The most abundant complex in both samples has a 1:1:1:1:1:1:1:1:1 stoichiometry, constituting 63.49% and 68.48% of the detected species in AncAmor Exo9 and AncOpis Exo9, respectively. Considering the number of subunits in each detected species reveals that 76.91% of subunits formed the correctly assembled Exo9 complex in AncAmor Exo9 and 72.09% of subunits formed the correctly assembled Exo9 complex in AncOpis Exo9.

| Species | AncAmor Exo9 | | AncOpis Exo9 | |
|---|---|---|---|---|
| | Mass/ Da | Abundance/ % | Mass/ Da | Abundance/ % |
| 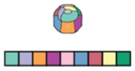 | 259916 | 63.49 | 260160 | 68.48 |
| | 260096* | | 260342* | |
| | 260288** | | 260518** | |
| 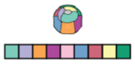 | — | — | 234113 | 9.76 |
| | — | | 234297* | |
| 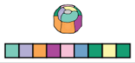 | 259544 | 3.54 | — | — |
| | 259726* | | — | |
| 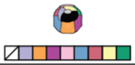 | 224613 | 5.12 | — | — |
| | 224792* | | — | |
| 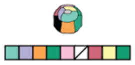 | 225198 | 7.69 | — | — |
| | 225371* | | — | |
| 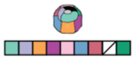 | — | — | 257781 | 16.91 |
| | — | | 257958* | |
| 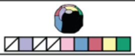 | — | — | 166918 | 1.49 |
| 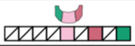 | — | — | 81763 | 1.58 |
| 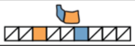 | 57959 | 17.10 | — | — |
|  | — | — | 27693 | — |
|  | — | — | 24023 | 1.78 |
|  | 30974 | 3.06 | — | — |

### Supplementary Table S3. Cryo-EM data collection, refinement and validation statistics

| | AncAmor Exo9 consensus | AncAmor Exo9 RNA-bound | AncAmor Exo9 RNA-unbound | AncAmor Opis9 | AncAmor Exo10 |
|---|---|---|---|---|---|
| **Data collection and processing** | | | | | |
| Magnification | 105,000 | 105,000 | 105,000 | 105,000 | 105,000 |
| Voltage (kV) | 300 | 300 | 300 | 300 | 300 |
| Electron exposure (e–/Å$^2$) | 50 | 50 | 50 | 50 | 50 |
| Defocus range (μm) | 1.0-2.5 | 1.0-2.5 | 1.0-2.5 | 1.0-2.5 | 1.0-2.5 |
| Pixel size (Å) | 0.831 | 0.831 | 0.831 | 0.831 | 0.831 |
| Symmetry imposed | C1 | C1 | C1 | C1 | C1 |
| Initial particle images (no.) | 2,627,464 | 2,627,464 | 2,627,464 | 715,355 | 1,376,574 |
| Final particle images (no.) | 998,128 | 30,171 | 502,163 | 82,930 | 43,485 |
| Map resolution (Å) | 2.7 | 3.6 | 2.8 | 3.7 | 3.6 |
| FSC threshold | 0.143 | 0.143 | 0.143 | 0.143 | 0.143 |
| **Refinement** | | | | | |
| R.m.s. deviations | | | | | |
| Bond lengths (Å) | 0.004 | 0.003 | 0.004 | 0.004 | 0.005 |
| Bond angles (°) | 0.458 | 0.541 | 0.736 | 0.798 | 0.780 |
| Validation | | | | | |
| MolProbity score | 1.38 | 2.16 | 1.81 | 1.85 | 2.06 |
| Clashscore | 6.94 | 10.8 | 10.42 | 12.97 | 14.43 |
| Poor rotamers (%) | 0.28 | 3.98 | 1.38 | 1.68 | 1.75 |
| Ramachandran plot | | | | | |
| Favored (%) | 98.30 | 97.15 | 97.06 | 97.75 | 96.69 |
| Allowed (%) | 1.70 | 2.85 | 2.94 | 2.25 | 3.28 |
| Disallowed (%) | 0.00 | 0.00 | 0.00 | 0.00 | 0.03 |

**Supplementary Table 4 | RNA oligonucleotides used for biochemical and structural analysis.**

| Name | Used for | Figure | Sequence (5‘-3‘) | Comment |
|---|---|---|---|---|
| RNA-36nts | RA | 2a,c, 3b, ED8a,b | [FAM] AUU AUU UAU UUA UUA AUU AUU UAU AUU UUA UUU AUU | |
| RNA-59nts | RA MP EM | 3c,d,e 4a,d 5a,c | [FAM] AUU CUA UCC CAG CGU CGU AUC UAU CCA AAA UUA AAC AAU AAU CAA UUA CAG UCC CUU UA | |
| RNA-18nts | RA MP EM | 3e, 4a,d, 5a,b,c | GAU ACG ACG CUG GGA UAG | Hybridization with RNA-59nts |
| RNA-80nts | RA | 5c | [FAM]AUU CUA UCC CAG CGU CGU AUC AUC AUU CUA UUC UAA ACC UAU UAU CCA AAA UUA AAC AAU AAU CAA UUA CAG UCC CUU UA | |
| RNA-35nts | MP | 3e | [FAM]AUU CUA UCC CAG CGU CGU AUC UAU CCA AAA UUA AA | |
| RNA-30nts | MP | 3e | [FAM]AUU CUA UCC CAG CGU CGU AUC UAU CCA AAA | |
| RNA-25nts | MP | 3e | [FAM]AUU CUA UCC CAG CGU CGU AUC UAUC | |
| RNA-35nts-2 | MP | ED9f | AUU CUA UCC CAG CGU CGU AUC UAU CCA AAA UUA AA | |
| RNA-21-nts | MP | ED9f | GAU ACG ACG CUG GGA UAG AAU | Hybridization with RNA-35nts-2 |

RA – RNase Assay; MP – Mass Photometry; EM – Cryo-Electron Microscopy;

ED – Extended Data

**Supplementary References**

1. Geoghegan, K. F. *et al.* Spontaneous α-N-6-Phosphogluconoylation of a "His Tag" inEscherichia coli:The Cause of Extra Mass of 258 or 178 Da in Fusion Proteins. *Anal. Biochem.* **267**, 169–184 (1999).
2. Marty, M. T. *et al.* Bayesian Deconvolution of Mass and Ion Mobility Spectra: From Binary Interactions to Polydisperse Ensembles. *Anal. Chem.* **87**, 4370–4376 (2015).
